# Modular and Distinct Plexin-A4/FARP2/Rac1 Signaling Controls Dendrite Morphogenesis

**DOI:** 10.1101/2020.01.15.908434

**Authors:** Victor Danelon, Ron Goldner, Edward Martinez, Irena Gokhman, Kimberly Wang, Avraham Yaron, Tracy S. Tran

## Abstract

Diverse neuronal populations with distinct cellular morphologies coordinate the complex function of the nervous system. Establishment of distinct neuronal morphologies critically depends on signaling pathways that control axonal and dendritic development. The Sema3A-Nrp1/PlxnA4 signaling pathway promotes cortical neuron basal dendrite arborization but also repels axons. However, the downstream signaling components underlying these disparate functions of Sema3A signaling are unclear. Using the novel *PlxnA4^KRK-AAA^* knock-in male and female mice, generated by CRISPR/cas9, we show here that the KRK motif in the PlxnA4 cytoplasmic domain is required for Sema3A-mediated cortical neuron dendritic elaboration but is dispensable for inhibitory axon guidance. The RhoGEF FARP2, which binds to the KRK motif, shows identical functional specificity as the KRK motif in the PlxnA4 receptor. We find that Sema3A activates the small GTPase Rac1, and that Rac1 activity is required for dendrite elaboration but not axon growth cone collapse. This work identifies a novel Sema3A-Nrp1/PlxnA4/FARP2/Rac1 signaling pathway that specifically controls dendritic morphogenesis but is dispensable for repulsive guidance events. Overall, our results demonstrate that the divergent signaling output from multifunctional receptor complexes critically depends on distinct signaling motifs, highlighting the modular nature of guidance cue receptors and its potential to regulate diverse cellular responses.

**Significance Statement:** The proper formation of axonal and dendritic morphologies is crucial for the precise wiring of the nervous system that ultimately leads to the generation of complex functions in an organism. The Semaphorin3A-Neuropilin1/Plexin-A4 signaling pathway has been shown to have multiple key roles in neurodevelopment, from axon repulsion to dendrite elaboration. This study demonstrates that three specific amino acids, the KRK motif within the Plexin-A4 receptor cytoplasmic domain, are required to coordinate the downstream signaling molecules to promote Sema3A-mediated cortical neuron dendritic elaboration, but not inhibitory axon guidance. Our results unravel a novel Semaphorin3A-Plexin-A4 downstream signaling pathway and shed light on how the disparate functions of axon guidance and dendritic morphogenesis are accomplished by the same extracellular ligand *in vivo.*

## Introduction

The development of precise neuronal connections critically depends on proper extension of axons and elaboration of dendrites. Studies of neurite outgrowth and axon guidance are well established (Barnes and Polleux, 2009; Kolodkin and Tessier-Lavigne, 2011; Stoeckli, 2018); and while numerous studies were conducted in invertebrates (Jan and Jan, 2010), much less is known about the signaling mechanisms controlling dendritic morphogenesis in the mammalian nervous system. Interestingly, molecular cues initially described as axonal guidance cues also control later events in dendrite development, suggesting that these cues are multifunctional (Whitford et al., 2002; Yaron et al., 2005; Tran et al., 2009; Smith et al., 2012; Mlechkovich et al., 2014; Nagel et al., 2015; Anzo et al., 2017).

The Semaphorin family of multifunctional cues has been mainly studied in the context of axon guidance (Jin and Strittmatter, 1997; Zanata et al., 2002; Huber et al., 2003; Toyofuku et al., 2005; Tong et al., 2007; Tran et al., 2007). Semaphorins can repel or attract axons by engaging different coreceptors (Castellani et al., 2000; Chauvet et al., 2007) or interacting with molecules in the local environment (Kantor et al., 2004). Interestingly, the Class 3 secreted Semaphorin-3A (Sema3A) can repel axons and promote dendrite branching in different neuronal populations both *in vitro* and *in vivo* (Gu et al., 2003; Fenstermaker et al., 2004; Yaron et al., 2005; Tran et al., 2009; Mlechkovich et al., 2014). However, the intracellular events orchestrating these divergent outputs, elicited by the same Sema3A cue, are largely elusive.

Sema3A signaling requires the binding to its obligate partner Neuropilin-1 (Nrp1) and formation of a complex with one of the Type-A Plexins (Plxns) (Tran et al., 2007). However, little was known about the specific Plxn signaling receptor involved in dendrite development until we showed the reduced dendritic complexity of *PlxnA4* KO deep cortical neurons in vivo (Tran et al., 2009) that phenocopied those observed in the *Nrp1^sema^*-mutants (Gu et al., 2003). We demonstrated *in vitro* that the ability of Sema3A to mediate growth cone collapse in DRG neurons versus dendritic arborization of cortical neurons is embedded in a modular nature of the PlxnA4 receptor, within its distinct cytoplasmic domains (Mlechkovich et al., 2014). We showed that the KRK motif in PlxnA4 can associate with the RhoGEF FARP2 and is required for dendrite elaboration *in vitro*. However, the downstream signaling elements controlling dendrite elaboration versus axon guidance remained elusive.

Regulators of small GTPases (GEFs and GAPs) may represent signaling-specificity elements for divergent Semaphorin functions. The RacGAP *β*2-Chimaerin is required for Sema3F-induced axonal pruning but not repulsion (Riccomagno et al., 2012). The PlxnA2 GAP activity is required for proper distribution of dentate gyrus granule cells, but not for mossy fiber-CA3 targeting in the developing hippocampus (Zhao et al., 2018). These studies, along with our work, suggest that Nrp/PlxnA complexes can recruit different GEFs and GAPs through distinct motifs, or the GAP domain itself, within the receptor to induce distinct cellular responses. Thus, we hypothesize that the KRK motif in PlxnA4 could serve as a recruitment site for specific GEFs, such as FARP2, to selectively confer Sema3A-dependent dendrite morphogenesis in cortical neurons *in vivo*.

Here, we generated a mouse line harboring an AAA substitution of the basic KRK motif in the PlxnA4 intracellular domain. Using this mouse line, we show that this conserved motif is required for Sema3A-induced dendritic elaboration in cortical neurons, but not for axonal repulsion or growth inhibition of several axonal tracts in the PNS and CNS. We demonstrate that cortical neuron dendrite development mediated by PlxnA4 signaling requires FARP2 *in vivo*. Furthermore, we show that Sema3A induces Rac1 activation in a FARP2-dependent manner in cortical neurons, and that this activity is selectively required for dendritic arborization, but not for growth cone collapse. Our work uncovers a distinct pathway for dendritic elaboration, specifically engaging the KRK motif of the multifunctional PlxnA4 receptor and its downstream signaling effectors.

## Materials and Methods

### Mouse strains

Mice harboring a point mutation in which the intracellular KRK (lysine-arginine-lysine) motif of Plexin-A4 was replaced by a triplet of alanines *(PlxnA4^KRK>AAA^)* were generated by CRISPR/Cas9-based homologous recombination of a 164 bp ssDNA oligonucleotide containing the KRK (5’-AAACGAAAA-3’) to AAA (5’-GCAGCAGCA-3’) mutation. A guide RNA (5’-CGCCATTGTCAGCATCGCGG-3’) was designed to target exon 20 of *PlxnA4* with minimal off-target effects. The ssDNA oligonucleotide had 76-79 bp homology arms around the KRK.AAA point mutation, with disruption of the TGG PAM sequence to TAG. *In vitro* transcribed Cas9 RNA (100ng/μl), the sgRNA (50ng/μl), together with the ssDNA (200ng/μl), were injected into one cell-fertilized embryos isolated from superovulated CB6F1 hybrid mice (F1 of BALB/ cAnNHsd inbred female x C57BL/6NHsd inbred male) mated with CB6F1 males from Harlan Biotech Israel. Injected embryos were transferred into the oviducts of pseudopregnant ICR females. Screening of tail tip DNAs was conducted by PCR genotyping and confirmed by sequencing. Heterozygous males were backcrossed with females of CD1 (ICR) genetic background.

For generation of the *Farp2* KO mouse line, two CRISPR sgRNAs were designed flanking a 18,167 bp sequence in the *Farp2* locus on Chr1 (GRCm38-mm10), including the promoter, the first utr exon and the second exon (with ATG). One sgRNA (5’ -TCTTAAGGACTTATTG CCAA-3’, chromosome 1: 93511211-93511211) was targeted to a region upstream of the *Farp2* gene promoter, and a second sgRNA (5’ – GTGAGGGCCTCATTCCGAAA-3’, chromosome 1: 93529378-93529397) to intron 2. The sgRNAs were designed using several CRISPR designing tools, and optimized for the best guides, including the following: the MIT CRISPR design tool (Hsu et al., 2013) and sgRNA Designer, Rule Sets 1 and 2 (Doench et al., 2014, 2016), in both the original sites and later in the Benchling implementations (www.benchling.com), SSC (Xu et al., 2015), and sgRNA scorer (Chari et al., 2015), in their websites. *In vitro* transcribed Cas9 RNA (100ng/μl) and the sgRNAs (50ng/μl) were injected into one cell-fertilized embryos isolated from superovulated CB6F1 hybrid mice (F1 of BALB/cAnNHsd inbred female x C57BL/6NHsd inbred male) mated with CB6F1 males from Harlan Biotech Israel. Injected embryos were transferred into the oviducts of pseudopregnant ICR females. Screening of tail tip DNAs was conducted by PCR genotyping and RT-PCR (see Fig. 1). Heterozygous males were backcrossed with females of CD1(ICR) genetic background.

**Figure 1.**
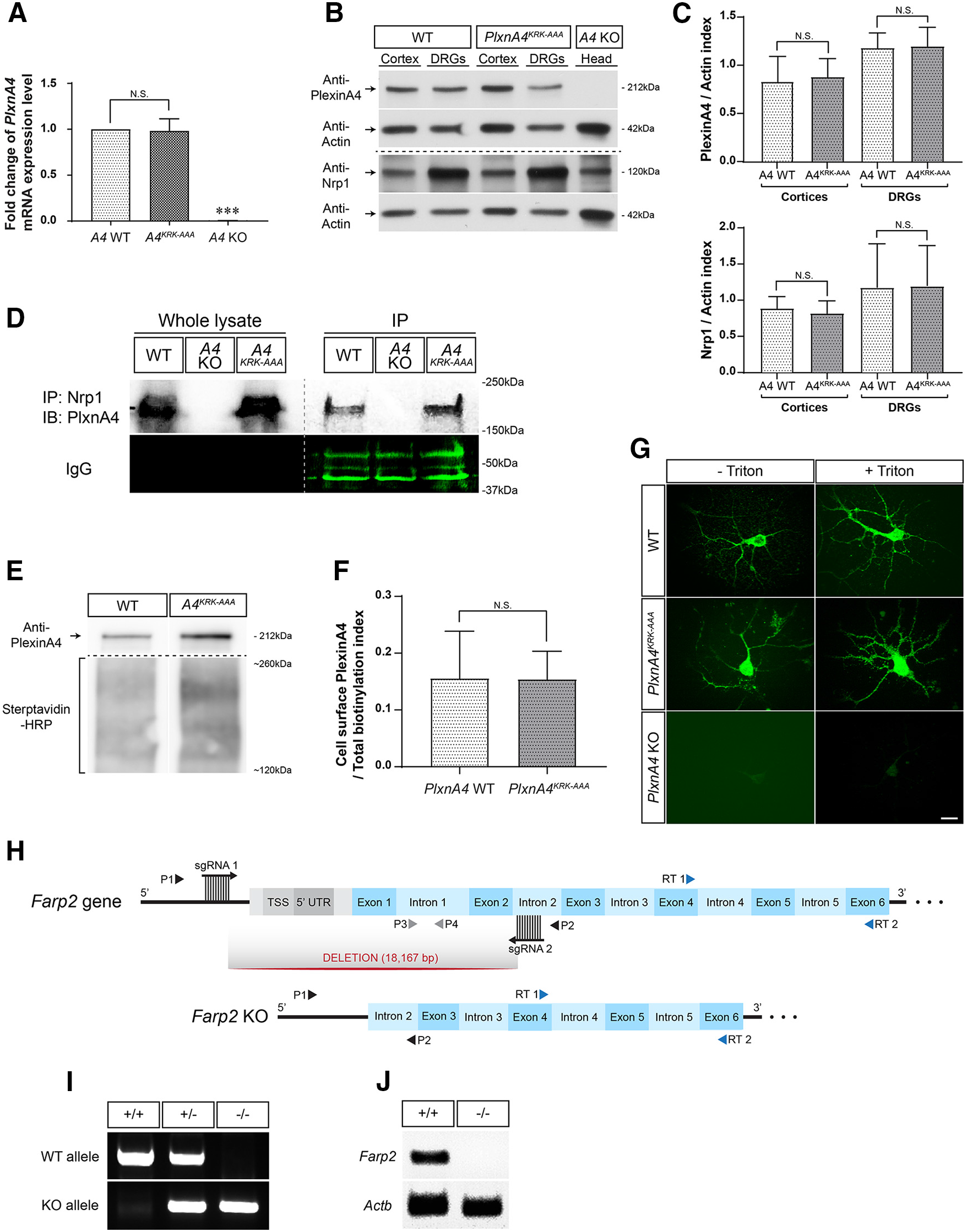
Generation of the *PlxnA4^KRA-AAA^* mutant mouse line and the *Farp2* KO mouse line. ***A***, qRT-PCR analysis revealed unchanged *PlxnA4* mRNA transcript levels in E13.5 *PlxnA4^KRK-AAA^* heads compared with WT littermate controls. *PlxnA4* KO heads were processed as well as a negative control. Quantification was performed using a relative quantification analysis (ΔΔC_t_) and presented as mean ± SEM. ***p <0.001 (Student’s *t* test). ***B, C**,* Western blot analysis of E13.5 DRGs and cortices extracted from *PlxnA4^KRK-AAA^* and WT littermate controls revealed similar Plexin-A4 and Nrp1 protein levels in both genotypes. Heads of E13.5 *PlxnA4* KO were used as a negative control. Data are mean ± SEM. Statistical analysis: Student’s *t* test. ***D**,* PlxnA4^KRK-AAA^, WT littermates, and *PlxnA4* KO cortices were subjected to anti-Nrp1 immunoprecipitation and probed using a Plexin-A4 antibody. No change in Nrp1-Plexin-A4 association was detected in the *PlxnA4^KRK-AAA^* mutant compared with WT. ***E, F**,* 7 DIV cortical neurons from E13.5 *PlxnA4^KRK-AAA^* and WT littermate embryos were subjected to a cell-surface biotinylation assay, followed by Western blot probing for Plexin-A4. The total mass of biotinylated proteins in the range of ~120-260 kDa, detected and quantified using streptavidin-HRP, was used as loading control. Data are mean ± SEM; N.S., Non Significant. ***G**,* Primary cortical neurons of *PlxnA4^KRK-AAA^,* WT littermates, and *PlxnA4* KO embryos were immunostained in culture using a Plexin-A4 antibody. Plexin-A4 protein levels and its spatial pattern in the *PlxnA4^KRK-AAA^* mutant were comparable with the WT, whereas virtually no Plexin-A4 was detected in the negative control KO. Scale bar, 25 μm. ***H***, Schematic representation of the *Farp2* locus on chromosome 1 (GRCm38-mm10) and the CRISPR-Cas9 KO design. Two sgRNAs flanking an 18,167 bp sequence were used: sgRNA 1 targeted a region upstream of the *Farp2* gene promoter, and sgRNA 2 targeted a sequence in intron 2. P1 and P2 (black triangles) represent primers used for detection of the KO allele in a PCR analysis of genomic DNA, targeting sequences outside of the deletion area (upstream of sgRNA1 and downstream of sgRNA2, respectively). Similarly, P3 and P4 (gray triangles) represent primers used for detection of the WT allele and were targeted at intron 1. RT 1 and 2 (blue triangles) represent primers used for reverse transcription analysis of *Farp2* gene expression levels; RT1 targeted a sequence comprised of the 3’ of exon 4 and 5’ of exon 5, and RT 2 targeted exon 6.1, PCR analysis of tail genomic DNA obtained from three E13.5 embryos of a *Farp2* Hetx Het cross, using primers 1-4 (P1-P4). Lengths of the WT and KO alleles are 551 and 271 bp, respectively. +/+, WT; +/−, *Farp2* heterozygous; —/—, *Farp2* KO. *J,* Reverse transcription analysis of WT (+/+) and KO (—/−) cDNA using either *Farp2* primers RT1-2 (top), or Actin *β* (Actb) primers as internal control (bottom). ***G, H***, Successful KO of the *Farp2* gene and its transcripts.

To investigate the role of FARP2 in mediating Sema3A cortical dendritic arborization, we used *Farp2* KO mice generated by Takegahara et al. (2010). The following oligonucleotide primers were used to identify the rearranged *Farp2* locus: primer 1 (5-ATCAAACTCCACCCTGA GGTCCATG-3), primer 2 (5-TTTGTAAACTGCAGCGTTTCTCTTC-3), and primer 3 (5-CTTCTGAGGGGATCGGCAAT-3).

Mice harboring the Thy1-EGFP (M line) transgene were obtained at The Jackson Laboratory (stock #007788 Tg-Thy1-EGFP-MJrs/J) and maintained by breeding with female C57BL/6J WT mice. *PlxnA4* KO mice were previously described (Yaron et al., 2005). Embryonic day (E) 0.5 was considered to be the day a vaginal plug was found. Newborn pups were weaned at postnatal day (P) 21 and housed under a 12 h light/ 12 h dark cycle at constant temperature with food and water *ad libitum*. All animal work in this study has been approved by the Institutional Animal Care and Use Committees Rutgers University Newark and the Weizmann Institute of Science.

### Primary neuron cultures

For the growth cone collapse assay, DRGs from E13.5 embryos were dissected and plated as explants in chambers coated with 10 μg/ml poly-D-ly-sine (Sigma Millipore, catalog #P6407) and 10 μg/ml laminin (Sigma Millipore, catalog #L2020). Growing media were as follows: Neurobasal-A (Thermo Fisher Scientific, catalog #10888-022) supplemented with 2% B-27 (Invitrogen, catalog #17504044), 1% penicillin-streptomycin (Biological Industries, catalog #03-031-1B), 1% L-glutamine (Biological Industries, catalog #03-020-1B), and 25 ng/ml NGF (Alomone Labs, catalog #AN-240). After 48 h, explants were treated with 0.1, 0.5, or 1 nm AP-Sema3A for 30 min, then fixed with 4% PFA 1 15% sucrose solution, and stained with phalloidin-rhodamine (Sigma Millipore, catalog #P1951, 1:250) for visualization of F-actin filopodia. Growth cones with no or few filopodia were considered as collapsed. Percentage of collapsed growth cones was calculated for each treatment group and control.

For the collagen axonal repulsion assay, DRGs from E13.5 embryos were dissected and placed in droplets of 2 mg/ml collagen (Roche Diagnostic, catalog#11179179001) together with COS1 aggregates transfected with either secreted Myc-Sema3A or control PAY1-GFP. Cocultures were grown for 48 h in the aforementioned Neurobasal-A growth medium + 12.5 ng/ml NGF, then fixed with 4% PFA, and stained with anti-tubulin *β*III antibody (R&D systems, Tuj1 clone, 1:1000). Extent of axonal repulsion was measured by the ratio between the lengths of the proximal axons (i.e., axons that are close to the COS1 aggregate) to the length of the distal axons.

Cortical neurons were obtained from E13.5 WT, or from *PlxnA4* KO, *PlxnA4^KRK-AAA^* and *Farp2* KO mice as previously described (Mlechkovich et al., 2014). When the experiments required, dissociated neurons were transfected with Farp2-HA-tagged (Mlechkovich et al., 2014) using Amaxa Mouse Neuron Nucleofector Kit catalog #VPG-1001. The neurons were plated onto glass coverslips (coated with poly-D-lysine, 0.1mg/ml, catalog #P6407) in 12-well plates (TPP 92412) and cultured with Neurobasal growth medium supplemented with 2% B-27 (catalog #17 504-044), 1% penicillin/streptomycin (catalog #15140122), and 1% Glutamax (catalog #35050) for 4-5 d.

### Production of AP-Sema3A

Alkaline phosphatase (AP)-tagged Sema3A was produced in HEK293T cells as previously described (Mlechkovich et al., 2014). Briefly, HEK293T cells (ATCC catalog #CRL-3216, RRID:CVCL_0063) were grown in DMEM supplemented with 10% FBS (VWR, catalog #97068-085) and 1% penicillin-streptomycin. The cells were transfected using BioT (catalog #B01-01) with an AP-Sema3A expression vector (Huber et al., 2005). The conditioned medium was concentrated using Amicon ultra-15 UFC 910024 (Sigma Millipore). AP-Sema3A concentration was determined by AP activity assay (using AP substrate buffer para-nitro-phenyl phosphate, Sigma Millipore) (Mlechkovich et al., 2014).

### Rac1 activity assay

Rac1 activation was measured by using a Rac1 activation assay kit (Rac1 Pull-down Activation Assay Biochem Kit bead pulldown format, Cytoskeleton, catalog #BK035), following the manufacturer’s protocol, with minor modifications. Briefly, fresh lysates of primary cortical neurons at DIV 5 incubated either with AP (5 nm) or Sema3A-AP (5 nm) for 30 min were incubated with the glutathione S-transferase (GST)-fused p21-binding domain of PAK1 (GST-PAK1-PBD) at 4°C for 1 h with gentle shaking, followed by pelleting of the PAK1-PBD beads by centrifugation at 3-5000 x *g* at 4°C for 1 min. After one wash with wash buffer, the beads were eluted in reduced 2x SDS sample buffer (Laemmli buffer) and analyzed by Western blot. The Rac1 antibody used was anti-Rac1 (Abcam, catalog #ab33186), concentration 0.6 μg/ml.

### Rac inhibitor EHT1864

For cortical neurons, EHT1864 (Tocris Bioscience, catalog #3872, batch #1J/197009), inhibitor of Rac family GTPases (Shutes et al., 2007; Onesto et al., 2008), was prepared following manufacturer guidelines. Cortical neurons were incubated with EHT1864 at different concentrations (2.5, 5, and 10 *μ*m) (Gaitanos et al., 2016) for 2 h at 37°C before treatment and also added with either 5 nm of AP-Sema3A or AP alone treatment. DRG explants were incubated with the specified EHT1864 concentration for 30 min at 37°C before Sema3A bath application. The same volume of H_2_O was added to the no inhibitor treatment control cells.

### Rac1 siRNA

The siRNA was purchased from Dharmacon. Rac1 siRNA: catalog #M-041170-01-0005, siGenome Mouse Rac1 (19 353) siRNA-SMART pol, 5 nmol. Target sequences: D-041170-01, GGACGAAGCUUGAUCUU AG; D-041170-02, AGACGGAGCUGUUGGUAAA; D-041170-03, GAU CGGUGCUGUCAAAUAC; and D-041170-04, GCAAAGUGGUAUCC UGAAG.

Scrambled siRNA: catalog #D-001206-13-05, siGENOME Non-Targeting siRNA Pool #1, 5 nmol. Target sequences: UAGCGACUA AACACAUCAA, UAAGGCUAUGAAGAGAUAC, AUGUAUUGGCC UGUAUUAG, and AUGAACGUGAAUUGCUCAA. The Rac1 siRNA or the Scramble was transfected to cortical neurons using Amaxa Mouse Neuron Nucleofector Kit catalog #VPG-1001.

### Immunohistochemistry

After 4-5 DIV, cortical neurons were treated with 5 nm AP-Sema3A or AP alone for 24 h, fixed in 4% PFA at 4°C for 15 min, and processed for immunocytochemistry, as previously described (Mlechkovich et al., 2014). Primary antibodies used were as follows: monoclonal rabbit anti-MAP2 (1:1000, Cell Signaling Technology, 4542S) and monoclonal mouse anti-HA (1:500, Santa Cruz Biotechnology, F7 sc-7392). Secondary fluorescently conjugated antibodies were AlexaFluor-488 donkey anti-mouse IgG (1:500, Jackson ImmunoResearch Laboratories) and Cy3 donkey anti-rabbit (1:1000, Jackson ImmunoResearch Laboratories). Coverslips were mounted on microscope slides using Mowiol (Sigma Millipore, catalog #81381)/10% p-phenylenediamine (catalog #78460).

For cell surface expression of PlxnA4 and PlxnA4^KRK-AAA^ protein, nonpermeabilized primary cortical neurons were immunostained using 1:1000 rabbit monoclonal Plexin-A4 (C5D1).

For whole-mount Neurofilament staining, embryos at E12.5 were collected into 1 x PBS, washed twice 5 min each with 1 x PBS + 0.2% Triton X-100 (PBST), and fixed with 4% PFA for 3 h at room temperature. Embryos were washed three times x 10 min with PBST, dehydrated by incubation in increasing concentrations of methanol in PBSx 1 (10%, 30%, 50%, and 80%, 1 h each), and stored overnight at 4°C in bleaching solution made of 6% H_2_O_2_ and 80% methanol in 1x PBS. Embryos were then washed three times x 1 h with 80% methanol in 1 x PBS, rehydrated with decreasing methanol concentrations in PBSx 1 (50%, 30%, and 10%, 1 h each), washed three times x 30 min with PBST, and blocked overnight at 4°C with 2% milk powder + PBST (PBSMT). Following 3 d incubation with primary mouse anti-Neurofilament antibody (2H3, DSHB, 1:200 in PBSMT) at 4°C, embryos were washed six times with PBSMT x 1 h and incubated overnight at 4° C with secondary HRP-coupled goat anti-mouse antibody (1:200) in PBSMT + 2% goat serum. Embryos were washed four times with PBSMT + 2% goat serum x 1 h, then two times with PBST x 1 h, and incubated for 30 min in 0.1 M Tris, pH 7.5, + DAB (Sigma Millipore, 1 tablet per 15 ml) prefiltered using a 0.45 μm filter. Development was performed by addition of H_2_O_2_ to the DAB solution (0.03% H_2_O_2_ final concentration), and stopped with two 1x PBS washes x 2 min, after fixation with 4% PFA for 15 min, and additional three 1 x PBS washes x 5 min.

For whole-mount TH staining, embryos at E13.5 were collected, and their sympathetic ganglia were exposed by dissection. Embryos were fixed with 4% PFA overnight at 4°C, washed with 1 x PBS + 1% Tween-20 (PBST) x 10 min, and incubated for 2 h in 0.9% NaCl made in 1x PBS. Dehydration was conducted as described for the anti-Neurofilament staining (but 15 min each instead of 1 h), followed by bleaching in 6% H_2_O_2_ + 20% DMSO in 80% methanol in 1 x PBS. Embryos were rehydrated (15min each), washed, blocked for 3 h at room temperature with 1x TBS, pH 7.4, + 1% Tween-20 (TBST) + 5% milk powder (TBSMT), then incubated for 2 d at 4°C in primary rabbit anti-TH antibody (Abcam, 1:200, in TBSMT + 5% DMSO). After 6 washes x 1 h with TBST, embryos were incubated overnight at 4°C with secondary HRP-coupled goat anti-rabbit antibody (1:200) in TBST + 2% goat serum, then washed five times x 1 h with TBST, followed by 1 h wash with 1x TBS. Development was performed as described for the anti-Neurofilament staining.

### Golgi staining

Adult (2-3 months old) mouse brains were processed for Golgi staining as previously described (FD Rapid GolgiStain Kit, catalog #PK401, FD NeuroTechnologies) for 10d (Tran et al., 2009). After incubation, all brains were blocked and embedded in OCT embedding medium (Tissue-Tek, catalog #4583). Sagittal sections (100 *μm)* were cut with a cryostat (Microm International, HM 505E) and mounted on charged microscopes slides (Diamond white glass). Staining procedures were followed exactly as described (FD NeuroTechnologies). Only layer 5 pyramidal neurons from the medial somatosensory cortex were included in our analyses. Differential interference contrast z-stack images were taken with an Axio Examiner Z1, Yokogawa spinning disk microscope (Carl Zeiss).

### Confocal images

High-resolution confocal *z*-stack images of MAP2-immunostained neurons were taken using an Axio Examiner Z1, Yokogawa spinning disk microscope (Carl Zeiss) with an oil-immersion 63x (NA 1.4) objective (PlanApo, Carl Zeiss). High-resolution confocal z-stack images of GFP-positive cortical neurons from WT *Thy1-EGFP, PlxnA4^KRK-AAA^; Thy1-EGFP* and *PlxnA4^+/KRK-AAA^/Farp2* +/−; *Thy1-EGFP* animals were taken using an LSM510 META microscope (Carl Zeiss). Series of optical sections were acquired in the *z* axis at 1 *μ*m intervals through individual layer 5 pyramidal neurons. Maximum projections of fixed images were analyzed using Fiji (is just ImageJ) software.

### Quantification of dendritic arborization

Dendritic arborization was analyzed with the following parameters:

#### Sholl analysis

All Golgi-labeled *z*-stack images were reconstructed using Fiji and Adobe Photoshop CS6, and all analyses were performed using the Fiji Sholl Analysis plugin (Ferreira et al., 2014). For cortical basal dendrite Sholl analysis, the starting radius is 10 *μ*m and the ending radius is 60 *μ*m from the center of the neuron soma; the interval between consecutive radii is 5 *μ*m (Mlechkovich et al., 2014).

#### Total dendritic length

The total dendritic length was measured using the ImageJ plugin NeuronJ (http://www.imagescience.org/meijering/software/neuronj/), calculated in microns (*μ*m).

#### Dendritic complexity index (DCI)

Dendritic order is defined as follows: primary dendrites were traced from the cell soma to the tip of the entire dendritic length, and secondary and tertiary dendrites were traced from the tip to the dendritic branch point using Adobe Photoshop software. Dendrites that were <10 *μ*m from the cell soma were disregarded. The DCI was calculated using the following formula: (∑branch tip orders + # branch tips)/(# primary dendrites) x (total arbor length) (Lom and Cohen-Cory, 1999; Peng and Tran, 2017).

For each parameter analyzed, at least 30-40 neurons were measured per condition, from three or four independent cultures/mice.

### Western blot

For Figure 1*B,C*, tissues were lysed with RIPA buffer (50 mm Tris, pH 7.4, 159 mm NaCl, 1% NP40, 0.1% SDS, 0.5% deoxycholate, 1 mm EDTA in ddH_2_O) supplemented with 1:25 cOmplete Protease Inhibitor Cocktail (Roche Diagnostic, catalog #11697498001) and 1:200 PMSF. Samples then underwent two cycles of 10-min-long incubation on ice followed by 10 s of vortexing. After 14,000 x *g* centrifugation for 15 min at 4°C, supernatant was collected and protein concentration was measured using Pierce BCA Protein Assay Kit (Thermo Fisher Scientific, catalog #23227) and Gen5 3.04 software (BioTek). Sample buffer X5 (125 mm Tris HCl, pH 6.8, 25% *β* -mercaptoethanol, 43.5% glycerol, 10% SDS, 0.05% bromophenol blue) was added to the samples that were further denatured by boiling at 95°C for 5min. Samples were separated by SDS-PAGE and electrophoretically transferred onto nitrocellulose membranes. Nonspecific antigen sites were blocked using 5% milk in TBS (10 mm Tris, pH 8, 150 mm NaCl) and 0.1% Tween-20 (TBST) for 1 h at room temperature. Antigen detection was performed by overnight incubation with appropriate antibodies: 1:1000 rabbit monoclonal Plexin-A4 (catalog #C5D1) anti-mouse (Cell Signaling Technology, catalog #3816), 1:1000 rabbit monoclonal Nrp1 (catalog #D62C6) anti-mouse (Cell Signaling Technology, catalog #3725), and 1:30,000 mouse anti-actin antibody C4 (MP Biomedicals, catalog #08691001). Antibodies were diluted in 5% BSA in TBST + 0.05% sodium azide. Membranes were then washed and incubated for 1 h in at room temperature in goat antirabbit HRP (Jackson ImmunoResearch Laboratories, 111-035-003) or goat anti-mouse (Jackson ImmunoResearch Laboratories, 115-035-003) secondary antibodies, following incubation in WesternBright HRP-substrate enhanced chemiluminescence (Advansta, catalog #K-12 045-D20) and exposure to Super RX-N X-Ray films (Fujifilm, catalog #47 410). Films were scanned and band intensities were quantified using ImageJ (Fiji) software.

For Figure 1D, mice were sacrificed using a CO_2_ chamber. The cortices were immediately removed from the skull and homogenized in RIPA buffer containing protease inhibitors (Roche cOmplete Ref. 11697498001). Homogenates were cleared by centrifugation at 500 x *g* twice for 7 min, and protein concentration was determined using the Bradford protocol (Bradford, 1976); samples were then boiled in gel-loading buffer and separated using SDS-PAGE, 10% for PlxnA4 (catalog #C5D1, Cell Signaling Technology), and IgG. Proteins were transferred to nitrocellulose membranes (Bio-Rad, catalog #1620097) and blocked with 5% nonfat milk in TBS with 0.05% Tween, for 1 h at room temperature. The membranes were incubated with primary antibodies (overnight at 4°C), washed, and reincubated with the secondary HRP-conjugated antibody (1:2000 dilution for anti-rabbit, GE Healthcare, NA934, lot #6969611; 1:2000 anti-mouse GENXA931; 1 h at room temperature). Peroxidase activity was visualized by using enhanced chemiluminescence kit (catalog #32106, Thermo Fisher Scientific). Enhanced chemiluminescence signal was exposed to BioBlue Lite films XR8813. Membranes were reprobed with a monoclonal antibody against IgG for control of protein loading. The images were quantified using Fiji software.

### Coimmunoprecipitation assay

Mice were sacrificed using a CO_2_ chamber, and the cortices were immediately removed from the skull and homogenized in RIPA-modified buffer (TBS 1x, 10% glycerol, 1% Triton X-100, and 1% NP40) supplemented with protease inhibitors. Briefly, the homogenates were centrifuged at 14,000 x *g* for 10 min, and 500 *μ*g of total protein from the supernatants was precleared with Protein G Sepharose 4 Fast Flow (GE Healthcare, catalog #GE17-0618-01) for 2 h at 4°C. After centrifugation, the precleared supernatants were incubated with anti-Nrp1 (R&D Systems, catalog #AF566) at 4°C for overnight. Then 10 *μ*l protein Protein G Sepharose 4 Fast Flow was added and incubated at 4°C for 4 h. The immunoprecipitates were washed with ice-cold RIPA-modified buffer, eluted with SDS sample buffer, and analyzed by Western blot. Membranes where then probed with anti-PlxnA4 (catalog #C5D1, Cell Signaling Technology) antibody. As a protein loading control, we first incubated the membranes with the secondary antibody anti-goat alone (primary antibody from goat).

### Cell-surface biotinylation assay

Cortical neurons from E13.5 embryos were extracted and grown in culture. At 7 DIV, a cell surface biotinylation assay was performed using a commercially available kit (Pierce Cell Surface Protein Isolation Kit, Thermo Fisher Scientific, catalog #89881). Eluates were subjected to immunoblot analysis using a Plexin-A4 antibody (Cell Signaling Technology, catalog #3816), as described above in Western blot. Cell lysates obtained following the biotinylation process were used as loading controls, and the total mass of proteins between ~120 and 260 kDa was used for quantification and normalization purposes.

### Real-time RT-PCR

Heads of E13.5 embryos were collected and subjected to RNA extraction. Briefly, heads were homogenized in TRI reagent (Sigma Millipore) and chloroform was added. Samples were mixed thoroughly, incubated at room temperature for 10 min, centrifuged at 13,000 rpm for 15 min at 4° C, and upper RNA-containing phase was collected. Precipitation of RNA was performed using isopropanol, after which samples were incubated at room temperature for 10 min, and centrifuged at 13,000 rpm for 15 min at 4°C. Cold 75% ethanol was added to the RNA pellet, and samples were stored overnight at — 80°C. The next day, ethanol was decanted, and samples were incubated at 65°C for 5 min to remove all traces of ethanol. RNA pellet was liquidized with 30 μl ultrapure DNase/RNase free H_2_O, and RNA concentration was measured using NanoDrop spectrophotometer. cDNA was then synthesized from 1 μl/RNA sample using an iScript kit (Bio-Rad, catalog #1708891), and *PlxnA4* expression levels were measured by qRT-PCR reaction using KAPA SYBR FAST ABI prism qPCR kit (Kapa Biosystems, catalog #KK4605). Primers (5’3’) were as follows: *PlxnA4,* GCAAGAACTTGATCCCGCCT and GTCACGGTGCATGGTTTCTC; *Actb*, GGCTGTATTCCCCTCCA TCG and CCAGTTGGTAACAATGCCATGT. Results were subjected to relative quantification analysis (ΔΔC_t_) using Microsoft Excel software.

### Experimental design and statistical analyses

For sequencing analysis of exon 20 in the *PlxnA4* locus experiments, a total of 3 animals were used. For PlxnA4 mRNA qRT-PCR analysis, a total of 6 animals/genotype were used, and a Student’s *t* test was performed. For Western blot analysis of Plexin-A4, Nrp1, and actin proteins in E13.5 DRGs and cortices extracted from *PlxnA4^KRK-AAA^,* WT littermate, and *PlxnA4* KO, a total of 4 mice/genotype were used from three independent experiments. For Western blot analysis of Nrp1:Plexin-A4 interaction, cerebral cortices were extracted from *PlxnA4^KRK-AAA^*, WT littermate, and *PlxnA4* KO, a total of 3 animals/genotype were used from three independent experiments. For analysis of the expression pattern of the PlxnA4 protein, primary cortical neurons obtained from WT, *PlxnA4^KRK-AAA^*, and *PlxnA4* KO embryos were immunostained using a Plexin-A4 antibody and a total of three independent cultures/genotypes. For analysis of Plexin-A4’s cell-surface expression level, three independent cell-surface biotinylation experiments were conducted and followed by an immunoblot analysis using a Plexin-A4 antibody. For the analysis of Sema3A-mediated growth cone collapse using DRG explants obtained from WT, *PlxnA4^KRK-AAA^,* and *Farp2* KO embryos, a total of at least 300 growth cones/group from three independent experiments were analyzed, and two-way ANOVA with *post hoc* Tukey test was performed. For the Sema3A-collagen axonal repulsion assay using DRG explants obtained from WT, *PlxnA4^KRK-AAA^,* and *Farp2* KO embryos, a total of at least 8 DRGs/genotype from three independent experiments were used, and a Student’s *t* test was performed. For Sema3A-treatment in primary cortical neuron culture obtained from WT, *PlxnA4* KO, *PlxnA4^KRK-AAA^*, and *Farp2* KO, a total of three independent cultures/genotype were used, and a Student’s *t* test and a two-way ANOVA followed by *post hoc* Tukey test were performed. For Golgi-stained layer V cortical neurons obtained from WT, *PlxnA4* KO, *PlxnA4^KRK-AAA^*, *Farp2* KO, and *PlxnA4*^+*/KRK-AAA*^/ *Farp2^+/−^*, a total of 3 animals/genotype were used. For analysis of layer V cortical neurons in WT, *PlxnA4^KRK-AAA^,* and in *PlxnA4^1/KRK-AAA^/ Farp2^+/−^,* using Thy1GFP mice, a total of 3 animals/genotype were used. For quantification of dendritic morphologies using Sholl analysis, dendritic length, DCI, and number of dendritic tips, one-way ANOVA and *post hoc* Tukey test were performed. For the analysis of the pattern of cutaneous sensory axons projections in WT, *PlxnA4^KRK-AAA^*, *Farp2* KO, or *PlxnA4* KO E12.5 embryos, using a Neurofilament antibody, a total of at least eight embryos/genotype from at least three independent experiments were used. For the analysis of sympathetic axon projections in WT, *PlxnA4^KRK-AAA^*, *Farp2* KO, or *PlxnA4* KO E13.5 embryos, a total of at least 10 embryos/genotype from at least three independent experiments were used. For the analysis of the anterior commissure in WT, *PlxnA4^KRK-AAA^*, *Farp2* KO, or *PlxnA4* KO, using hematoxylin staining, a total of at least seven brains/genotype from at least three independent experiments were used. For the analysis of the Rac signaling in Sema3A-dependent growth cone collapse of phalloidin-rhodamine-stained DRG neurons obtained from WT and *PlxnA4^KRK-AAA^* DRGs, a total of at least 300 growth cones/group from three independent experiments were analyzed, and a two-way ANOVA with *post hoc* Tukey test was performed. For the analysis of the Rac signaling inhibition (using EHT) in Sema3A-dependent dendritic growth in WT cortical primary neurons, a total of three independent cultures were used, and a two-way ANOVA followed by *post hoc* Tukey test was performed. For the analysis of Sema3A-dependent dendritic arborization in the *Farp2* KO with Rac inhibitor, three independent cultures were performed, and a two-way ANOVA followed by *post hoc* Tukey test was performed. For the Western blot of Rac1 knockdown experiments, using a specific Rac1 siRNA, three independent primary WT neuron cultures were used, and a Student’s *t* test was performed. For Sema3A treatment in the Rac1 knockdown experiments, a total of three independent primary WT cortical neuron cultures were used, and two-way ANOVA followed by *post hoc* Tukey test was performed. For the pulldown assay showing the effect of 30 min stimulation of Sema3A on Rac1 activation, three independent WT, *PlxnA4^KRK-AAA^*, and *Farp2* KO primary neuron cultures were used, and Student’s *t* test and one-way ANOVA were performed. For DRG growth cone collapse and cortical neuron dendritic arborization quantifications, data obtained were analyzed using GraphPad Prism Software. For DRG axonal repulsion, data were analyzed using Microsoft Excel software. Data are shown as mean ± SEM.

## Results

### Generation of the *PlxnA4^KKK-AAA^* mutant and *Farp2* KO mouse

To explore the role of the KRK motif in the PlxnA4 cytoplasmic domain, we have generated by CRISPR the *PlxnA4^KRK-AAA^* line that harbors a KRK substitution to AAA in the endogenous PlxnA4 receptor. The modified genomic area (220 bp) from tail genomic DNA was sequenced, and only the desired modification was detected. The mutation had no effect on PlxnA4 mRNA or total protein levels, as shown using biochemical analysis (*Fig. 1A* and Fig. 1*B,C*, respectively). Moreover, we have not detected any change in the levels of the PlxnA4 coreceptor (Fig. 1*B,C*), nor change in the association of Nrp1 with PlxnA4 in the *PlxnA4^KRK-AAA^* mutants (Fig. 1D). Furthermore, the KRK to AAA substitution did not affect the cell surface expression of PlxnA4 on cortical neurons, as revealed by detection of the biotinylated receptor (Fig. 1*E,F*), or its dendritic localization (Fig. 1*G*). Overall, our analysis shows that the KRK to AAA mutation did not alter the expression and cellular distribution of Plexin-A4. In parallel, we have generated a *Farp2* KO mouse by CRISPR (Fig. 1*H*). Using PCR analysis of tail genomic DNA and reverse transcription analysis of progenies from the *Farp2* KO line, we also demonstrated successful deletion of the *Farp2* locus and its transcripts (Fig. 1*I,J*). For both mutant lines, the injected animals (F0) were crossed with WT CD1 mice three times. In all the experiments described below, WT littermate controls were used.

### DRG sensory axons from *PlxnA4^KRK-AAA^* and *Farp2* KO animals are sensitive to Sema3A inhibition and repulsion

The *PlxnA4^KRK-AAA^* and the *Farp2* mutant animals allowed us to systemically examine their role in Sema3A signaling using multiple *in vitro* assays. We first assessed the acute growth cone collapse assay. DRG axonal growth cones from E13.5 *PlxnA4^KRK-AAA^* homozygous mutant, grown for 2 DIV and visualized with phalloidin-rhodamine, were able to respond to Sema3A-mediated collapse in a dosage-dependent manner, similar to the WT control (Fig. 2*A,B*). In line with our results from the *PlxnA4^KRK-AAA^* mutant neurons, DRG axonal growth cones from mice deficient of FARP2 GEF (*Farp2* KO), which is a downstream target of the KRK motif in PlxnA4, collapsed at percentages comparable with the WT control following Sema3A treatment (Fig. 2*A,B*). Next, we tested the response to the prolonged Sema3A-mediated axonal repulsion in *PlxnA4^KRK-AAA^* and *Farp2* mutant animals. For this, we cocultured DRG explants with COS1 cell aggregates either transfected with GFP (control) or a construct expressing Sema3A. Both DRG sensory axons from *PlxnA4^KRK-AAA^* and *Farp2* KO explants were repelled from the Sema3A source, similar to WT DRG axons (Fig. 2*D*). Quantification of axonal repulsion using the proximal/distal axonal outgrowth ratio measurement as previously described (Yaron et al., 2005) showed no significant difference between *PlxnA4^KRK-AAA^* or *Farp2* KO explants compared with their respective littermate WT controls in response to Sema3A repulsion (Fig. 2*E*; *p* = 0.48). These results suggest that the KRK motif in the PlxnA4 receptor and its downstream target FARP2 are dispensable for both Sema3A-mediated acute growth cone collapse and prolonged axonal repulsion.

**Figure 2.**
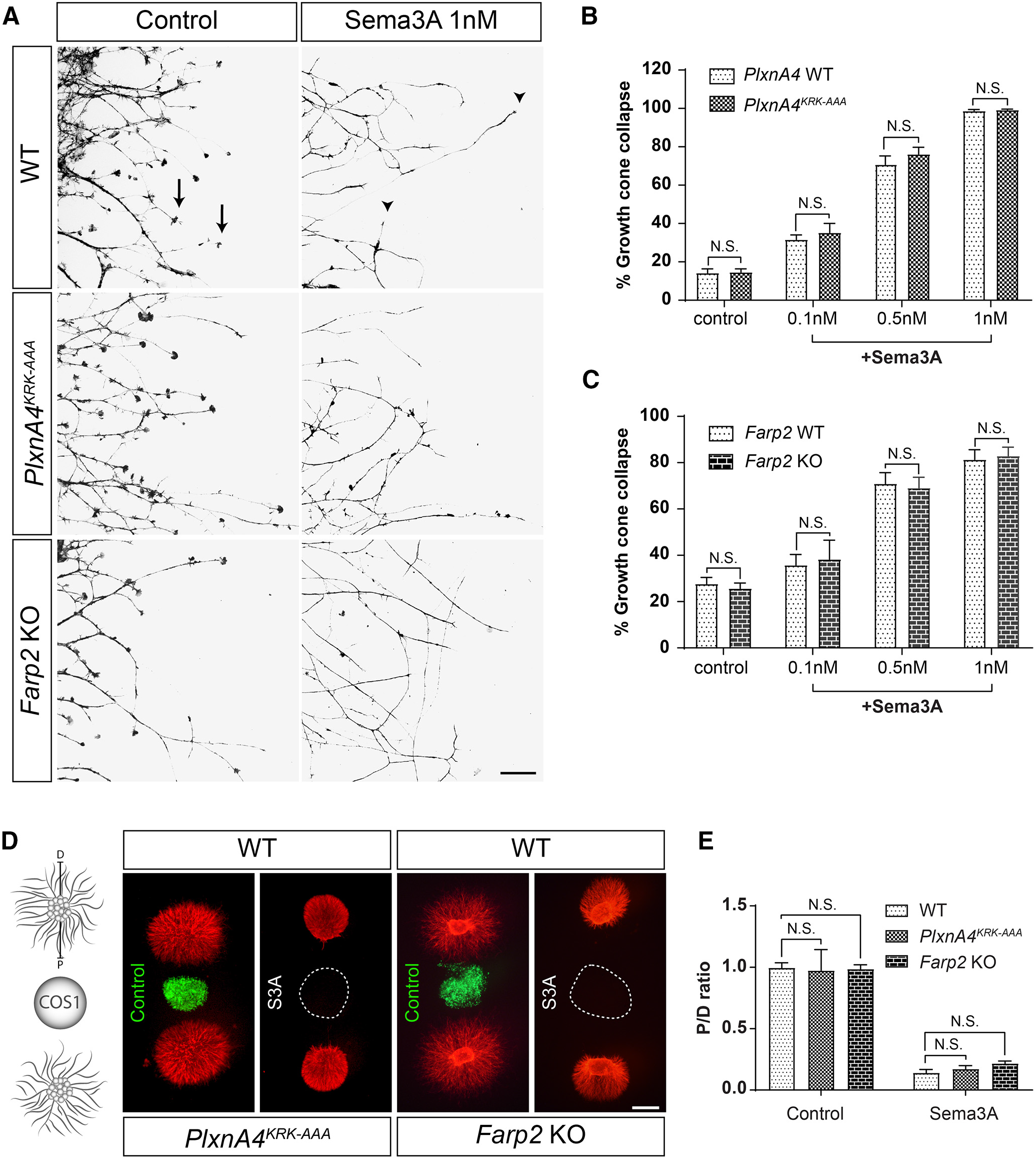
*PIxnA4^KRK-AAA^* and *Farp2* KO DRG axons show intact Sema3A-dependent responses *in vitro. **A**,* DRG explants from WT, *PIxnA4^KRK-AAA^*, and *Farp2* KO E13.5 embryos were grown for 48 h, treated with 0.1, 0.5, or 1 nm AP-Sema3A (only 1 nm is shown) or control conditioned media and stained using phalloidin-rhodamine to visualize growth cone collapse. Arrows indicate intact growth cones. Arrowheads indicate collapsed growth cones. ***B, C***, Quantification of collapse response as a mean percentage of collapsed growth cones out of the total ± SEM. Statistical analysis: two-way ANOVA with *post hoc* Tukey test. Scale bar, 50 μm. ***D***, DRG explants from E13.5 *PIxnA4^KRK-AAA^* and WT littermates or *Farp2* KO and WT littermates were cocultured in collagen droplets for 48 h with a COS1 aggregate either secreting myc-Sema3A (dashed circle) or expressing control PAY1-GFP (green). Cultures were visualized using anti-tubulin Class III immunostaining. ***E**,* Quantification of axonal repulsion using the proximal/distal (P/D) ratio, as indicated in the schematic representation of the collagen axonal repulsion assay in ***D***. Data are mean ± SEM. Statistical analysis: Student’s *t* test; N.S., Non Significant. Scale bar, 500 μm.

### Cortical neurons from *PlxnA4^KRK-AAA^* and *Farp2* KO animals are nonresponsive to Sema3A-induced dendrite elaboration

As *Sema3A* KO brains have previously been shown to display similar cortical neuron basal dendrite morphologic defects (Nakamura et al., 2017) similar to those observed in *PlxnA4* KO (Tran et al., 2009; Mlechkovich et al., 2014; Peng and Tran, 2017), we asked whether the KRK motif of PlxnA4 and its downstream target FARP2 are required for Sema3A-mediated cortical dendrite branching. Embryonic cortical neurons at E13.5 were dissociated from WT, *PlxnA4^KRK-AAA^* mutant, *PlxnA4* KO, and *Farp2* KO animals as previously described (Tran et al., 2009; Mlechkovich et al., 2014; Peng and Tran, 2017) and grown for 5 DIV followed by Sema3A treatment for 24 h and fixation for anti-MAP2 immunocytochemistry to visualize all dendrites (Fig. 3*A*). Using four different measurements, we quantified the number of dendritic intersections (by Sholl analysis; Fig. 3*B*), the total dendritic length (Fig. 3*C*), the DCI (Fig. 3*D*), and the average number of dendritic tips (Fig. 3*E*). Consistent with previous results, Sema3A-treated WT neurons showed increased number of dendrites compared with the control-treated (Student’s *t* test, Sholl: 10 μm: *t* = 4.775, *p* = 0.0088; 15 μm: *t* = 5.422, *p* = 0.0056; 20 μm: *t* = 5.706, *p* = 0.0047; 25 μm: *t* =5.119, *p* = 0.0069, 30μm: *t* = 4.443, *p* = 0.0113; 35μm: *t* = 4.458, *p* = 0.0112; 40μm: *t* =3.999, *p* = 0.0161; 45μm: *t* =3.934, *p* = 0.0170; 50 μm: *t* =3.807, *p* = 0.0190; 55 μm: *t* = 2.952, *p* = 0.0419; 60μm: *t* = 2.910, p = 0.0437; length: *t* =5.022, *p* = 0.0074; DCI: *t*= 7.175, *p* = 0.0020; and dendritic tips: *t*= 6.553, *p* = 0.0028). Next, we asked whether neurons obtained from *PlxnA4^KRK-AAA^, Farp2* KO, and *PlxnA4* KO show sensitivity to Sema3A treatment. We found that *PlxnA4^KRK-AAA^* mutant and *Farp2* KO neurons were insensitive to Sema3A stimulation, similarly to the *PlxnA4* KO (two-way ANOVA: Sholl: 10 μm: *p* = 0.0023, *F* =7.554; 15 μm: *p* < 0.0001, *F* =14.41; 20 μm: *p* < 0.0001, *F* =18.04; 25 μm: *p* < 0.0001, *F* =15.59; 30 μm: *p* = 0.0002, *F* = 12.09; 35 μm: *p* = 0.0001, *F* = 13.05; 40 μm: *p* = 0.0004, *F* =10.59; 45μm: *p* = 0.0004, *F* =10.72; 50μm: *p* = 0.0015, *F* = 8.284; 55 μm: *p* = 0.0422, *F* =3.437; length: p = 0.0001, *F* =13,64; DCI: *p* < 0.0001, *F* = 27; dendritic tips: *p* < 0.0001, *F* = 25.50). Similarly, there were no significant differences in dendritic morphology as measured by Sholl analysis, dendritic length, DCI, and average number of dendritic tips between AP-treated WT neurons versus AP or Sema3A-treated *PlxnA4^KRK-AAA^* mutant, *PlxnA4* KO, and *Farp2* KO neurons. In addition, we tested whether higher Sema3A concentration could induce dendrite growth in *PlxnA4^KRK-AAA^* mutant and *Farp2* KO neurons. *PlxnA4^KRK-AAA^* and *Farp2* KO cortical neurons were treated with 10 nM of Sema3A for 24 h. We analyzed the dendritic morphology measured by Sholl analysis, dendritic length, DCI, and dendritic tips. We found that both *PlxnA4^KRK-AAA^* and *Farp2* KO cortical neurons remained nonresponsive, even to the higher concentration of Sema3A (two-way ANOVA, Sholl: 10 μm: *p* = 0.0075, *F* = 7.565; 15 μm: *p* = 0.0002, *F* = 19.6; 20 μm: *p* < 0.0001, *F* = 22.55; 25 μm: p = 0.0001, *F* = 20.66; 30 μm: *p* = 0.0005, *F* =15.11; 35μm: *p* = 0.0006, *F* =14.54; 40μm: *p* = 0.0018, *F* = 11.25; 45 μm: *p* = 0.0016, *F* = 11.49; 50 μm: *p* = 0.0025, *F* =10.30; 55μm: *p* = 0.0085, *F* = 7.27; 60μm: *p* = 0.0132, *F* = 6.342; length: *p* = 0.0007, *F* = 13.92; DCI: *p* < 0.0001, *F* = 28.55; dendritic tips: *p* = 0.0002, *F* = 18.58) (Fig. 4). Therefore, in contrast to DRG sensory neurons, these results suggest that *PlxnA4^KRK-AAA^* mutant and *Farp2* KO cortical neurons are selectively nonresponsive to Sema3A-mediated dendrite elaboration.

**Figure 3.**
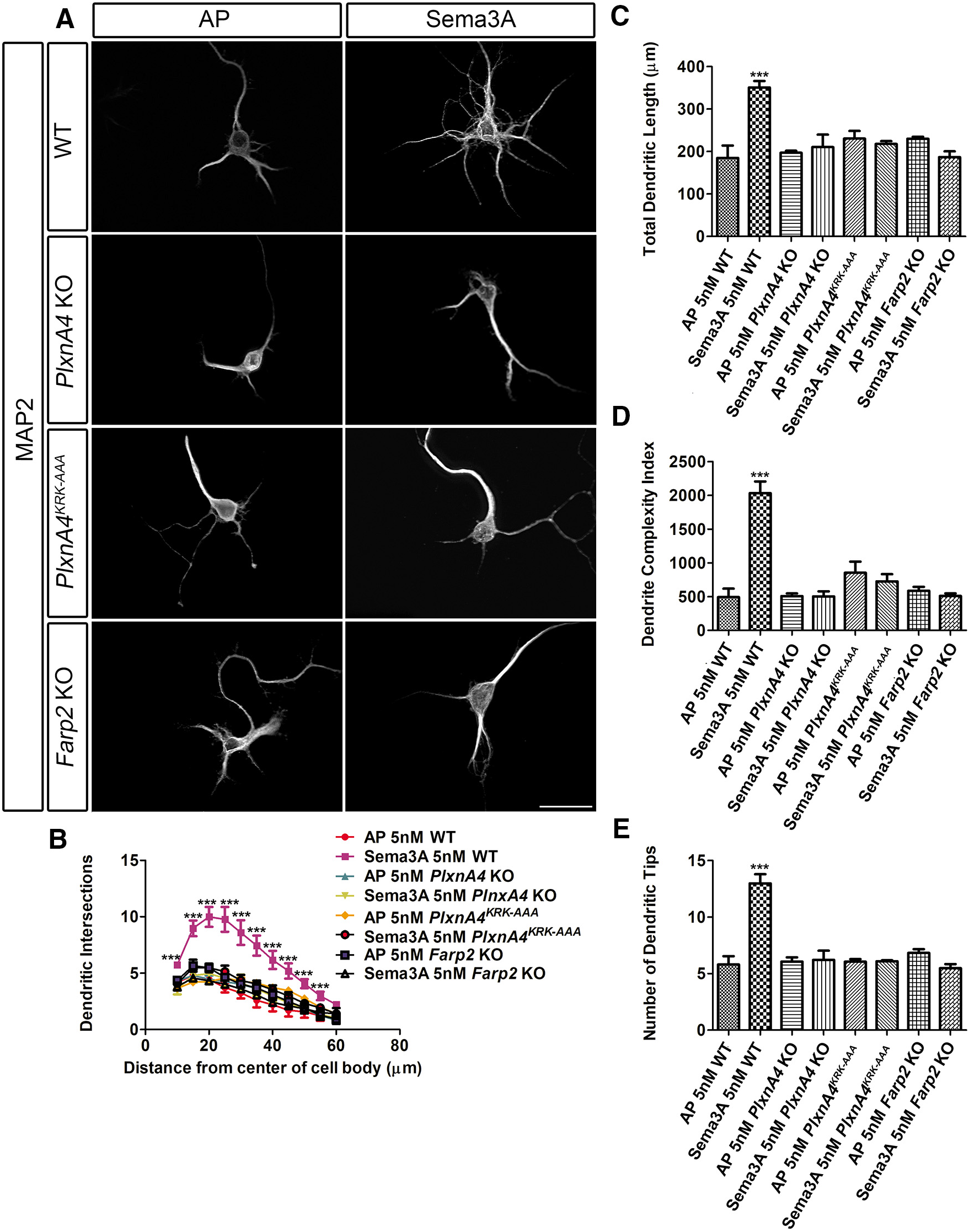
Both *PlxnA4^KRK-AAA^* and *Farp2* KO neurons are nonresponsive to Sema3A-induced cortical neuron dendrite outgrowth *in vitro. **A**,* Representative confocal micrographs of dissociated primary cortical neurons obtained from WT, *PlxnA4* KO, *PIxnA4^KRK-AAA^*, and *Farp2* KO E13.5 embryos. The neurons were treated with AP or Sema3A 5 nm for 24 h. ***B-E***, Sholl analysis quantified dendritic growth and branching in WT, *PlxnA4* KO, *PlxnA4^KRK-AAA^*, and *Farp2* KO cortical neurons (***B***), measuring the number of dendritic intersections, the total dendritic length (***C***), the DCI (***D***), and the number of dendritic tips (***E***). Data are mean ± SEM from three independent cultures of each genotype. ***p < 0.001 (two-way ANOVA with *post hoc* Tukey test). Scale bar, 25 μm.

**Figure 4.**
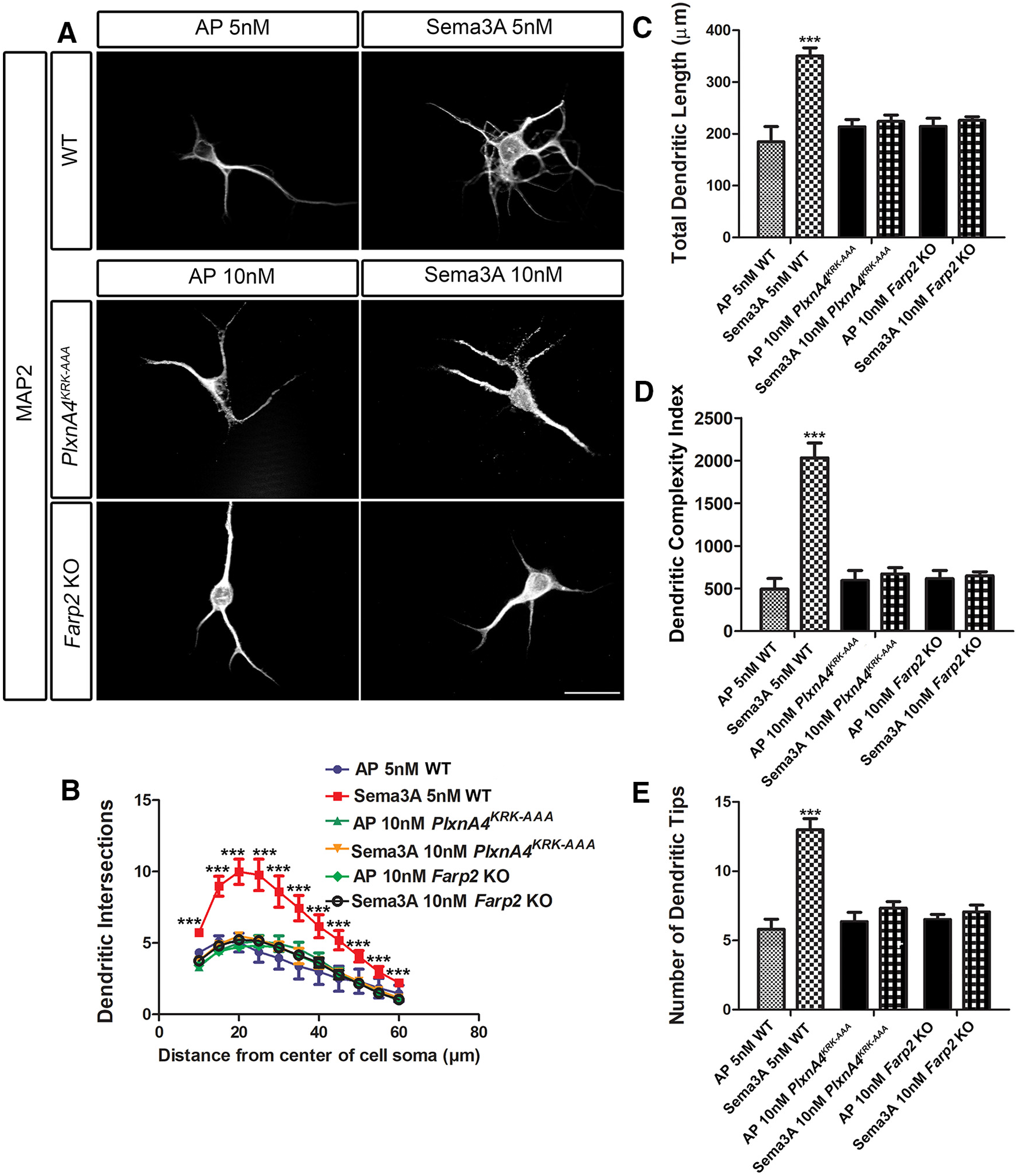
Both *PlxnA4^KRK-AAA^* and *Farp2* KO cortical neurons are nonresponsive, even to higher concentrations of Sema3A. ***A***, Representative confocal micrographs of dissociated primary cortical neurons obtained from WT, *PlxnA4^KRK-AAA^*, and *Farp2* KO E13.5 embryos. The WT neurons were treated with AP or Sema3A 5 nm, and the *PlxnA4^KRK-AAA^* and *Farp2* KO neurons were treated with AP or Sema3A 10 nm for 24 h. *B-E,* Sholl analysis quantified dendritic growth and branching in WT, *PlxnA4^KRK-AAA^,* and *Farp2* KO cortical neurons (***B***), measuring the number of dendritic intersections, the total dendritic length (***C***), the DCI (***D***), and the number of dendritic tips (***E***). Data are mean ± SEM from three independent cultures of each genotype. ***p < 0.001 (twoway ANOVA with *post hoc* Tukey test). Scale bar, 25 μm.

### *PlxnA4^KRK-AAA^* and *Farp2* KO animals exhibit reduced basal dendritic arborization in layer 5 cortical neurons, but not axonal repulsion defects *in vivo*

Next, we asked whether *PlxnA4^KRK-AAA^* homozygous mutant and *Farp2* KO embryos display dendritic elaboration phenotypes in layer 5 pyramidal neurons *in vivo*. We performed Golgi staining to uniformly label all dendritic processes in adult brains of WT, *PlxnA4^KRK-AAA^* mutant, *PlxnA4* KO, and *Farp2* KO animals. We observed severe reduction in the basal dendritic arbors of layer 5 cortical neurons in the *PlxnA4^KRK-AAA^* mutant, *PlxnA4* KO, and *Farp2* KO brains compared with the WT (Fig. 5*A*). As revealed by Sholl analysis, there was a significant decrease in dendritic intersections between WT and *PlxnA4^KRK-AAA^* mutant or *PlxnA4* KO or *Farp2* KO (Fig. 5*B*; Sholl: 15 μm: *p* < 0.0001, *F* =50.83; 20 μm: *p* < 0.0001, *F* =32.55; 25 μm: *p* < 0.0001, *F* = 23.39; 30 μm: *p* = 0.0001, *F* =18.50; 35 μm: *p* < 0.0001, *F* = 20.56; 40 μm: *p* = 0.0006, *F* =13.08; 45 μm: *p* = 0.0017, *F* =10.16; 50 μm: *p* = 0.0014, *F* = 10.67; 55 μm: *p* < 0.0003, *F* = 15.59; 60 μm: *p* = 0.0009, *F* = 11.79). In addition, the total dendritic length, the DCI, and average number of dendritic tips for *PlxnA4^KRK-AAA^* mutant, *PlxnA4* KO, and *Farp2* KO were lower compared with WT (Fig. 5*C-E*; length: *p* < 0.0001, *F* = 20.14; DCI: *p* = 0.0005, *F* =13.81; dendritic tips: *p* < 0.0001, *F* = 21.94). To provide genetic evidence that FARP2 is indeed downstream of PlxnA4 signaling, we generated compound heterozygous animals, *PlxnA4^KRK-AAA/+^; Farp2*^+/−^. The trans-heterozygous animals also exhibit reduced basal dendritic arbors, phenocopying the *PlxnA4* KO (Fig. 5*A-E*; Sholl analysis between WT and *PlxnA4^KRK-AAA/+^;Farp2^+/−^* and *PlxnA4* KO; Sholl: 15 μm: *p* = 0.0011, *F* =15.93; 20μm: *p* < 0.0001, *F* =38.17; 25 μm: *p* = 0.0002, *F* = 25.58; 30 μm: *p* = 0.0004, *F* = 20.52; 35 μm: *p* = 0.0003, *F* = 23.31; 40 μm: *p* = 0.0011, *F* = 16.07; 45 μm: *p* = 0.0023, *F* =12.83; 50 μm: *p* = 0.0016, *F* =14.43; 55 μm: *p* = 0.0002, *F* = 25.78; 60 μm: *p* = 0.0015, *F* =14.63; length: *p* = 0.0002; *F* = 25.71; DCI: *p* = 0.0005, *F* = 19.45; dendritic tips: *p* = 0.0003, *F* = 23.90). In addition, we compared the number of dendritic intersections, dendritic length, DCI, and number of dendritic tips between WT and *PlxnA4^KRK-AAA/+^* or *Farp2^+/−^* single heterozygous animals. There was not a significant difference between WT and *PlxnA4^KRK-AAA/+^* or *Farp2*^+/−^ heterozygous animals (images not shown); in contrast, there were significant differences between *PlxnA4^+/KRK-AAA^* single heterozygous or *Farp2^+/−^* single heterozygous and *PlxnA4^KRK-AAA^* mutant or *PlxnA4* KO or *Farp2* KO or compound heterozygous animals *PlxnA4^KRK-AAA/+^;Farp2^+/−^* (compared with *PlxnA4^+/KRK-AAA^* Sholl: 15 μm: *p* = 0.0156, *F* = 4.601; 20 μm: *p* = 0.0002, *F* =13.32; 25 μm: *p* < 0.0001, *F* =18; 30 μm: *p* < 0.0001, *F* =24.58; 35 μm: *p* = 0.0001, *F* = 14.49; 40 μm: *p* = 0.0013, *F* =8.544; 45 μm: *p* = 0.0033, *F* =6.923; 50 μm: *p* = 0.0027, *F* = 7.251; 55 μm: *p* = 0.0009, *F* =9.292; 60 μm: *p* = 0.0021, *F* =7.641; length: *p* < 0.0001, *F* = 20.18; DCI: *p* < 0.0001, *F* = 28.21; dendritic tips: *p* < 0.0001, *F* = 23.37; compared with *Farp2^+/−^* Sholl: 15 μm: *p* = 0.008, *F* =5.531; 20 μm: *p* = 0.0001, *F* =13.87; 25 μm: *p* < 0.0001, *F* =15.45; 30 μm: *p* < 0.0001, *F* =18.43; 35 μm: *p* = 0.0004, *F* =10.96; 40 μm: *p* = 0.0039, *F* = 6.625; 45 μm: *p* = 0.0065, *F*= 5.822; 50 μm: *p* = 0.0073, *F* =5.652; 55 μm: *p* = 0.0069, *F* =5.748; 60 μm: *p* = 0.0289; *F* =3.819; length: *p* < 0.0001, *F* =16.53; DCI: *p* < 0.0001, *F* = 37.28; dendritic tips: *p* < 0.0001, *F* =20.32).

**Figure 5.**
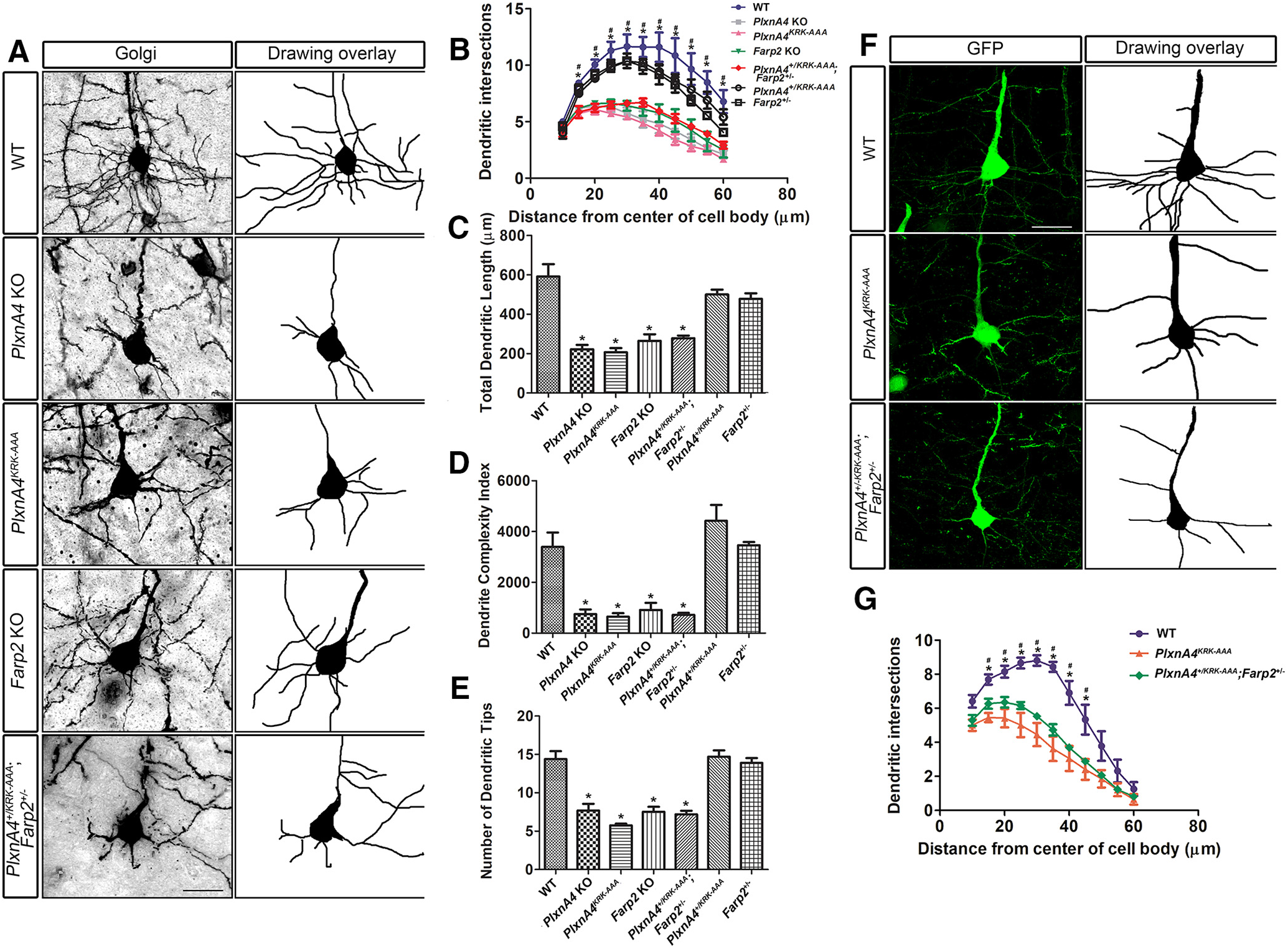
Reduced basal dendritic arborization in layer 5 cortical neurons from *PlxnA4^KRK-AAA^* and *Farp2* KO adult animals. ***A***, Golgi-stained images of brain (2-to 3-month-old) sections of WT, *PlxnA4* KO, *PlxnA4^KRK-AAA^, Farp2* KO, and *PlxnA4^+/KRK-AAA^/Farp2^+/−^*. ***B***, Sholl analysis quantification revealed reduced and altered branching patterns of basal dendrites from layer V cortical neurons in *PlxnA4* KO, *PlxnA4^KRK-AAA^, Farp2* KO, and *PlxnA4^+/KRK-AAA^/Farp2^+/−^* compared with WT and to *PlxnA4^+/KRK-AAA^* or *Farp2^+/−^* littermates. ***C-E**,* Quantifications of total dendritic length (***C***), DCI (***D***), and dendritic tips (***E***) in all genotype analyzed. Data are mean ± SEM. *p < 0.05 (one-way ANOVA followed by *post hoc* Tukey test). ***F, G***, Representative images of the layer 5 cortical neurons and Sholl analysis obtained from a WT, *PlxnA4^KRK-AAA^*, and *PlxnA4^+/KRK-AAA^/Farp2^+/−^* Thy1-GFP mice. Statistical analysis: one-way ANOVA followed by *post hoc* Tukey test. Data are mean ± SEM; *n* = 3 brains/genotype. #p < 0.05, statistical difference between WT and *PlxnA4^KRK-AAA^* mutant or *PlxnA4* KO or *Farp2* KO in ***B*** and ***G***. \**p* < 0.05, statistical difference between WT and *PlxnA4^KRK-AAA/+^/Farp2^+/−^* or *PlxnA4* KO in ***B*** and ***G***. Scale bars: ***A, F***, 25 μm.

While the Golgi staining is a great method for assessing overall neuronal morphologies, the impregnation of the silver nitrate is arbitrary for each neuron. Therefore, to confirm that indeed layer 5 neuron basal dendrites are exhibiting the reduced dendritic arbor phenotype observed, we crossed the *PlxnA4^KRK-AAA^* mutant or the *Farp2* KO with Thy1-EGFP reporter mice to genetically label layer 5 neurons. We used the M line of Thy1-EGFP to sparsely label specific layer 5 neurons for optimal visualization of the entire dendritic arbor of each neuron. Consistent with our Golgi staining analysis, we found that *PlxnA4^KRK-AAA^* mutant;Thy1-EGFP^+^ and *PlxnA4^KRK-AAA/+^;Farp2^+/-^;Thy1*-EGFP^1^ (trans-heterozygous) neurons displayed significantly reduced basal dendritic arbors compared with WT Thy1-EGFP^+^ neurons (Fig. 5*F,G*; Sholl: 15 μm: *p* < 0.0025, *F* =13.94; 20 μm: *p* < 0.0076, *F* =9.531; 25μm: *p* < 0.0031, *F* =12.99; 30 μm: *p* < 0.0006, *F* = 21.14; 35μm: *p* < 0.0007, *F* = 20.72; 40 μm: *p* = 0.0045, *F* = 11.44; 45 μm: *p* = 0.0177, *F* = 6.968).

In contrast to dendritic development, both *PlxnA4^KRK-AAA^* mutant and *Farp2* KO embryos at E12.5 showed normal sensory cutaneous axon patterning in the PNS, comparable with the WT, as opposed to the defasciculated, highly branched, and misguided axonal projections seen in the PlxnA4 null (Fig. 6*A*, arrows). Furthermore, it is well established that Sema3A-PlxnA4 signaling can exert axonal repulsion on different neuronal populations in both the PNS and CNS (Gu et al., 2003; Huber et al., 2005; Yaron et al., 2005). In addition to repelling peripheral DRG axons, PlxnA4 is also required in the PNS for the proper projections of sympathetic axons, and for the formation of the anterior commissure in the CNS (Yaron et al., 2005; Waimey et al., 2008). Therefore, we asked whether the KRK motif in the PlxnA4 receptor is dispensable only for PNS DRG axon repulsion or can this be a general principal for Sema3A signaling in axon guidance. TH staining of the sympathetic ganglia showed normal axonal patterns in *PlxnA4^KRK-AAA^* mutant and *Farp2* KO E13.5 embryos similar to WT control, whereas *PlxnA4* KO animals displayed disorganized axonal projections toward the midline (Fig. 6*B*). Hematoxylin staining of coronal sections from WT, *PlxnA4^KRK-AAA^*, *PlxnA4* KO, and *Farp2* KO brains at P30 showed normal anterior commissure projections in both *PlxnA4^KRK-AAA^* mutant and *Farp2* KO animals, similar to the WT, but in contrast to the missing commissure in the *PlxnA4* KO (Fig. 6*C*).

**Figure 6.**
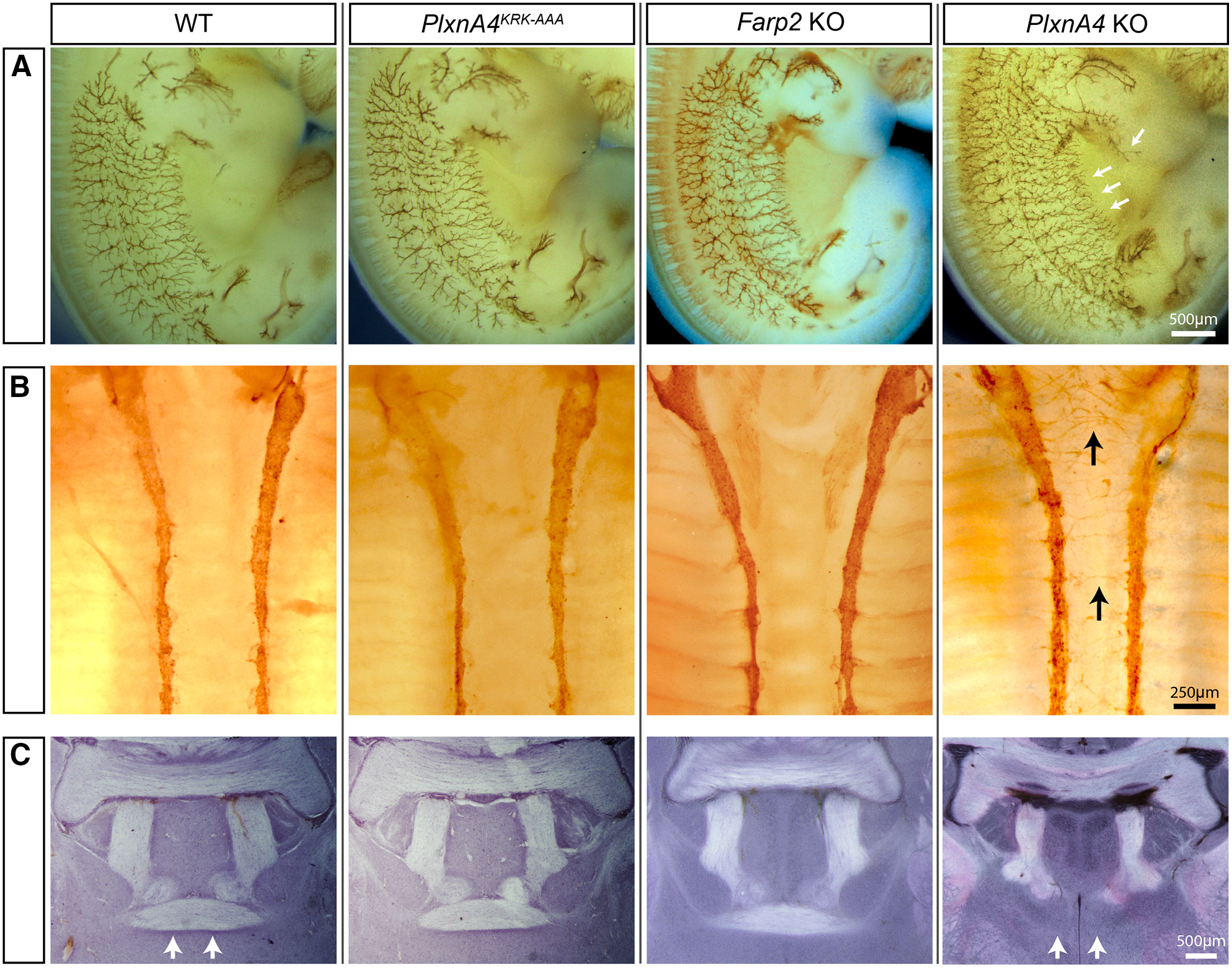
*PlxnA4^KRK-AAA^* and *Farp2* KO animals show normal axonal projections in the PNS and CNS *in vivo.* ***A***, WT, *PlxnA4^KRK-AAA^, Farp2* KO, or *PlxnA4* KO E12.5 embryos were immunostained using a Neurofilament antibody to visualize cutaneous sensory axons pattern. *PlxnA4^KRK-AAA^* and *Farp2* KO embryos were comparable with the WT, whereas the *PlxnA4* KO exhibited an expected hyperinnervation, including a previously described discrete phenotype (white arrows). Scale bar, 500 μm. ***B***, WT, *PlxnA4^KRK-AAA^, Farp2* KO, or *PlxnA4* KO E13.5 embryos were immunostained using a TH antibody to visualize sympathetic axons. *PlxnA4^KRK-AAA^* and *Farp2* KO embryos were comparable with the WT, whereas the *PlxnA4* KO exhibited an expected medial protrusion of sympathetic axons (black arrows). Scale bar, 250 μm. ***C***, WT, *PlxnA4^KRK-AAA^, Farp2* KO, or *PlxnA4* KO P30 brains were stained using hematoxylin. *PlxnA4^KRK-AAA^* and *Farp2* KO embryos were comparable with the WT, whereas the *PlxnA4* KO exhibited an expected disrupted formation of the anterior commissure (white arrows). Scale bar, 500 μm.

Together, these results suggest that the PlxnA4^KRK^-FARP2 signaling pathway is necessary to promote layer 5 pyramidal neuron basal dendrite elaboration but is dispensable for axonal repulsion *in vitro* and *in vivo*.

### Inhibition of Rac activity perturbs Sema3A-mediated cortical neuron dendrite elaboration, but not axon growth cone collapse

One of the major targets of FARP2 GEF is the small GTPase Rac1, but not other members of the Rho GTPase family: CDC42 or RhoA (Kubo et al., 2002; Toyofuku et al., 2005; He et al., 2013). A previous study showed that overexpression of FARP2 in murine primary neurons induced shortened neurites (Kubo et al., 2002); however, the physiological importance of FARP2-Rac1 signaling pathway in the nervous system is not clear. Therefore, we first asked whether Rac activation is required downstream of Sema3A-signaling for axonal growth cone collapse and dendrite morphogenesis using a pharmacological approach. We used a validated pan-Rac GTPase inhibitor, EHT1864 (Shutes et al., 2007; Onesto et al., 2008), in a dosage-dependent manner on cultured WT DRG neurons and found that this inhibitor had no effect on Sema3A-induced growth cone collapse (Fig. 7*A-C*). However, WT-dissociated cortical neurons treated with the Rac inhibitor for 2 h before Sema3A stimulation, at concentrations of 2.5, 5, or 10 *μ*m, showed complete inhibition of dendritic elaboration, compared with Sema3A treated neurons without the inhibitor. In contrast, there were no significant differences in dendritic morphology between AP-treated WT neurons versus AP-treated+EHT at different concentrations (2.5, 5, and 10 *μ*m) (Fig. 7*D*). Quantifications of dendritic intersections by Sholl analysis (Fig. 7*E*), total dendritic length (Fig. 7*F*), DCI (Fig. 7*G*), and average number of dendritic tips (Fig. 7*H*) showed significant differences between WT neurons treated with Sema3A only compared with Sema3A + 2.5 *μ*m EHT, Sema3A + 5 *μ*m EHT, Sema3A + 10 *μ*m EHT, 2.5, 5, or 10 *μ*m EHT alone or control (no Sema3A, no EHT) (two-way ANOVA, Sholl: 15 μm: *p* = 0.0019, *F* =7.822; 20 μm: *p* = 0.0012, *F* =8.652; 25 μm: *p* = 0.0005, *F* = 10.19, 30 μm: *p* = 0.0020, *F* =7.798.117; 35 μm: *p* = 0.0019, *F* = 7.895; 40 μm: *p* = 0.0181, *F* = 4.492; length: *p* = 0.0276, *F* = 3.674; DCI: *p* = 0.0030, *F* = 9.454; and dendritic tips: *p* = 0.0013, *F* =7.381). These results establish that Rac activation is required for Sema3A-mediated dendrite outgrowth and branching.

**Figure 7.**
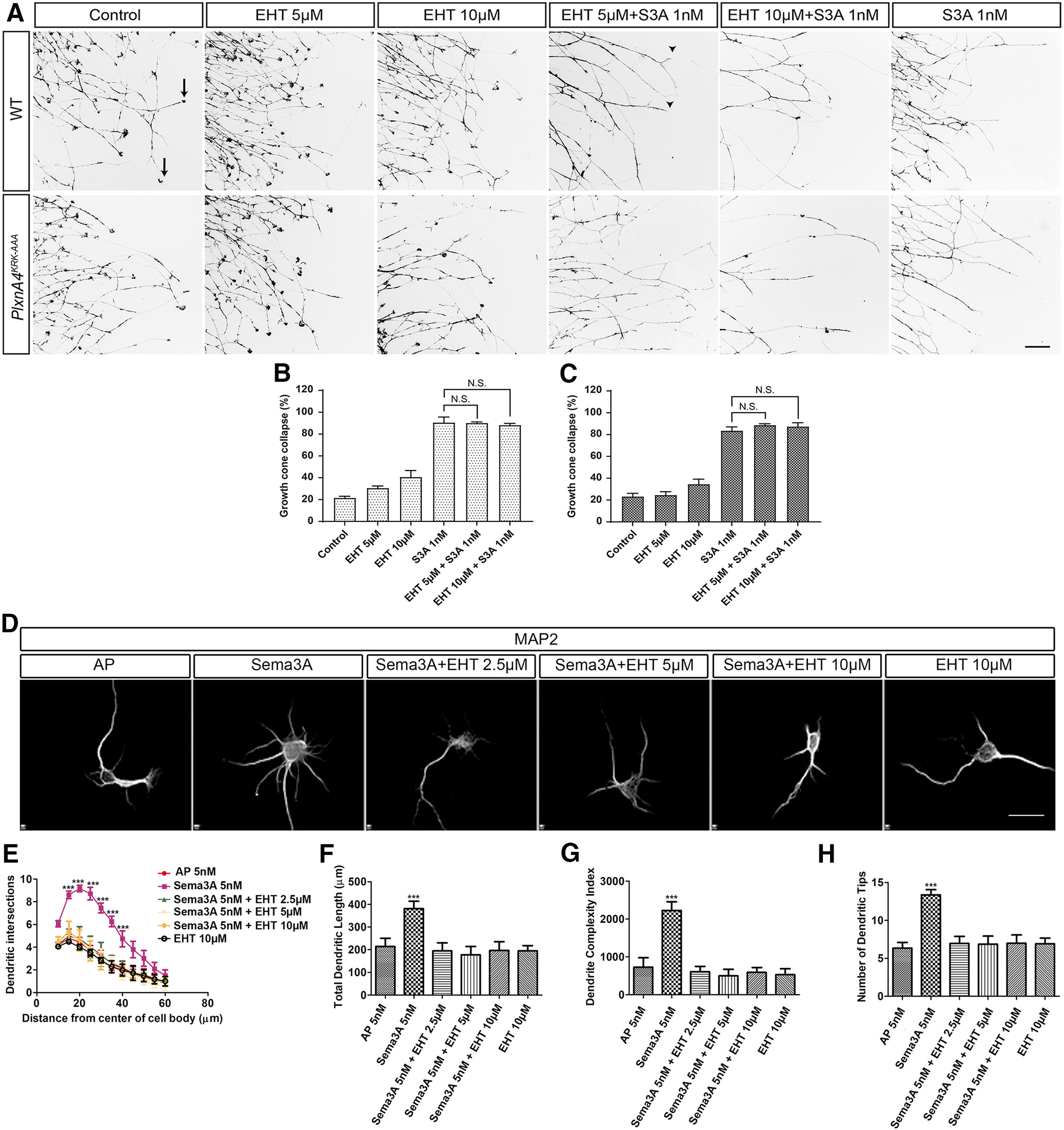
Inhibition of Rac signaling abolishes Sema3A-mediated cortical neuron dendrite elaboration but does not hinder Sema3A-dependent growth cone collapse of WT or *PlxnA4^KRK-AAA^* DRG axons *in vitro. **A**,* DRG explants from WT and *PlxnA4^KRK-AAA^* littermates E13.5 embryos were grown for 48 h, treated for 30 min with the pan-Rac inhibitor EHT 1864 at a concentration of 5 *μ*m, 10 *μ*m, or media alone as control. Then, 1 nm Sema3A or control conditioned media was added for 30 min, followed by fixation and phalloidin-rhodamine staining for assessment of growth cone collapse. Black arrows indicate intact growth cones. Arrowheads indicate collapsed growth cones. ***B, C***, Quantification of collapse response as a mean percentage of collapsed growth cones out of the total ± SEM in WT and *PlxnA4^KRK-AM^* axons, respectively; N.S., Non Significant. Scale bar, 50 μm. ***D***, Representative confocal micrographs of dissociated primary cortical neurons obtained from WT E13.5 embryos. The neurons were treated with 5 nm AP, 5 nm Sema3A, 5 nm Sema3A + 2.5 *μM* EHT, 5 nm Sema3A + 5 *μM* EHT, and 5 nm Sema3A + 10 *μM* EHT. ***E-H***, Sholl analysis of dendritic intersections (**E**), total dendritic length (***F***), the DCI (***G***), and number of dendritic tips (***H***) for all treatment conditions described above. Data are mean ± SEM from three independent cultures. ***p < 0.001 (two-way ANOVA with *post hoc* Tukey test). Scale bars: ***A, D***, 25 μm.

### Sema3A-mediated cortical neuron dendrite elaboration requires Rac1 activation by FARP2 GEF

To support our hypothesis that Sema3A-mediated dendrite elaboration involves a FARP2-Rac signaling pathway, we used *Farp2* KO cortical neurons, which we showed are insensitive to Sema3A-induced dendritic elaboration (Fig. 3). First, we demonstrate that by overexpressing *Farp2* cDNA into *Farp2* KO neurons, the sensitivity to Sema3A-induced dendritic elaboration is restored compared with *Farp2* KO neurons transfected with *Farp2* cDNA without Sema3A treatment (Fig. 8*A,B*; two-way ANOVA, Sholl: 10 μm: *p* = 0.0081, *F* =12.24; 15 μm: *p* = 0.0011, *F* = 24.47; 20 μm: *p* = 0.0007, *F* = 27.95; 25 μm: *p* = 0.0008, *F* = 26.81, 30 μm: *p* = 0.0086, *F* = 11.93). However, *Farp2* KO neurons were unable to elaborate dendrites when transfected with *Farp2* cDNA 1 2.5 *μ*m EHT and treated with Sema3A. Sholl analysis showed that the number of dendritic intersections significantly decreased between 10 and 30 μm away from the cell soma in *Farp2* KO neurons rescued with *Farp2* cDNA and treated with Sema3A 1 EHT compared with *Farp2* KO rescued with FARP2 cDNA and treated with Sema3A only (Fig. 8*A,B*; two-way ANOVA, Sholl: 10 μm: *p* = 0.0132, *F* =10.05; 15 μm: *p* = 0.0015, *F* = 22.08; 20 μm: *p* = 0.0023, *F* =19.24; 25 μm: *p* = 0.0021, *F* =19.84, 30 μm: *p* = 0.0021, *F* =19.87).

**Figure 8.**
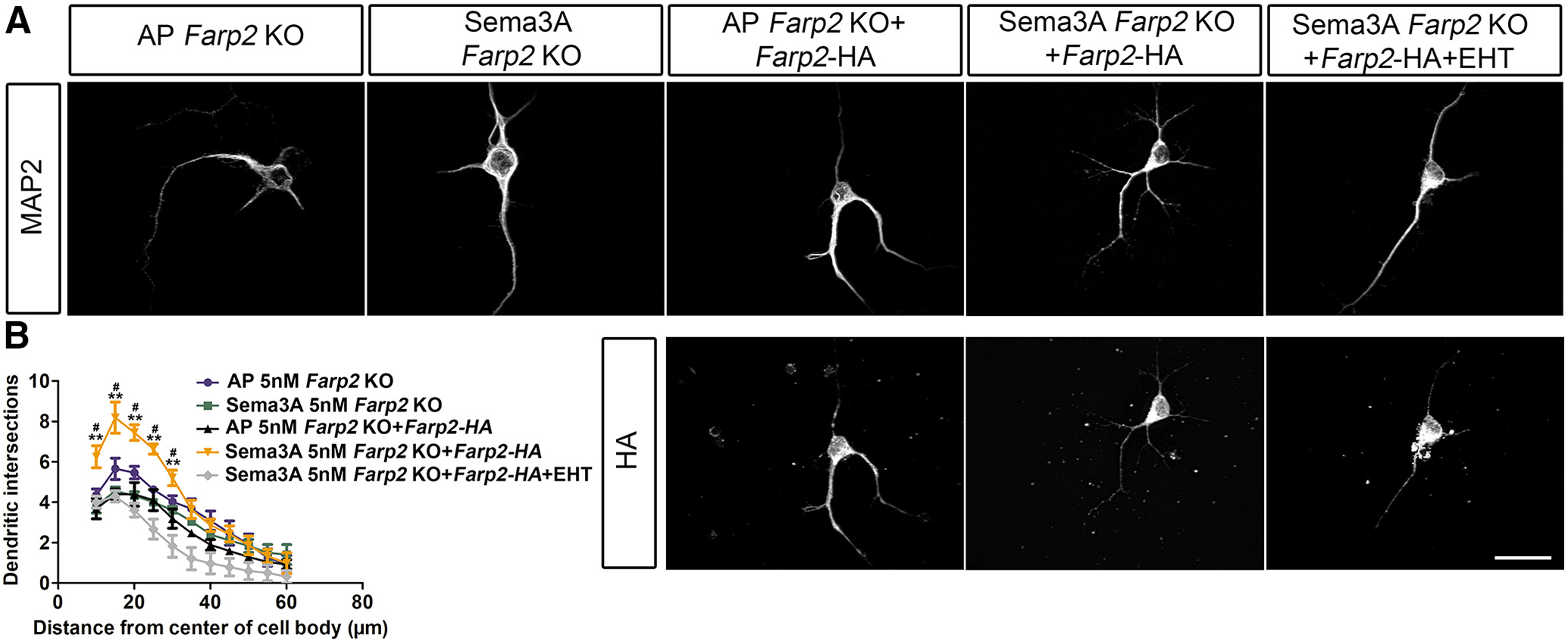
Sema3A-dependent dendritic arborization relies on the FARP2 GEF and downstream Rac1 GTPase activation. ***A***, Representative confocal micrographs of dissociated primary neurons obtained from *Farp2* KO animals. The neurons were treated with 5 nm AP, treated with 5 nm Sema3A, transfected with *Farp2-HA* cDNA and treated with 5 nm AP, transfected with *Farp2-HA* cDNA and treated with 5 nm Sema3A and transfected with *Farp2-HA* cDNA, treated with 5 nm Sema3A + 2.5 uM EHT for 24 h. ***B***, Sholl analysis of dendritic intersections in the five different groups. Data are mean ± SEM from three independent cultures. **, #p < 0.01 (two-way ANOVA with *post hoc* Tukey test). “Statistical difference between *Farp2* KO+*Farp2-HA* and *Farp2 KO+Farp2’* H’ I Sema3A. ‘Statistical difference between *Farp2 KO+Farp2’HA+Sema3A* and *Farp2 KO+Farp2’HA+Sema3A* + EHT. Scale bars: ***A***, 25 μm.

While the results from our pharmacological experiments are consistent with Rac1 as the downstream target of FARP2, EHT inhibits all Rac activity (Shutes et al., 2007; Onesto et al., 2008) activity. Using siRNA specific to Rac1, we found that WT cortical neurons transfected with Rac1 siRNA not only expressed a significantly lower level of Rac1 (Fig. 9*A,B*; Student’s *t* test, *t* =4.245, *p* = 0.0132), but were also nonresponsive to Sema3A-induced dendrite outgrowth and branching compared with the Sema3A-scramble siRNA (Fig. 9*C-G*; two-way ANOVA, Sholl: 15 μm: *p* = 0.0111, *F* =10.81; 20 μm: *p* = 0.0032, *F* =17.16; 25 μm: *p* = 0.0026, *F* =18.48; 30 μm: *p* = 0.0047, *F* =15.08; 35 μm: *p* = 0.0055, *F* =14.14; length: *p* < 0.0001, *F* =112.6; DCI: *p* < 0.0001, *F* = 70.5; dendritic tips: *p* = 0.0006, *F* = 29.49). Finally, we asked whether Rac1 activity is altered following Sema3A signaling in primary cortical neurons. Using a biochemical assay to pull down the downstream target of Rac1, PAK1, we quantified the level of active Rac1 in its GTP bound state (for details, see Materials and Methods). We found that the level of Rac1-GTP increased in WT primary cortical neurons following 30 min treatment with AP-tagged Sema3A compared with the vehicle control AP treatment (Student’s *t* test *t* = 3.168, *p* = 0.0339). Moreover, we found that Sema3A treatment on *PlxnA4^KRK-AAA^* and on *Farp2* KO cortical neurons did not induce any significant increase in Rac1 activity (Fig. 9*H,I*, one-way ANOVA *p* = 0.0022; *F* = 20.14). Together, our results demonstrate that Sema3A-induced cortical neuron dendrite elaboration requires the activation of the PlxnA4^KRK^ motif, which signals through FARP2 GEF to activate Rac1.

**Figure 9.**
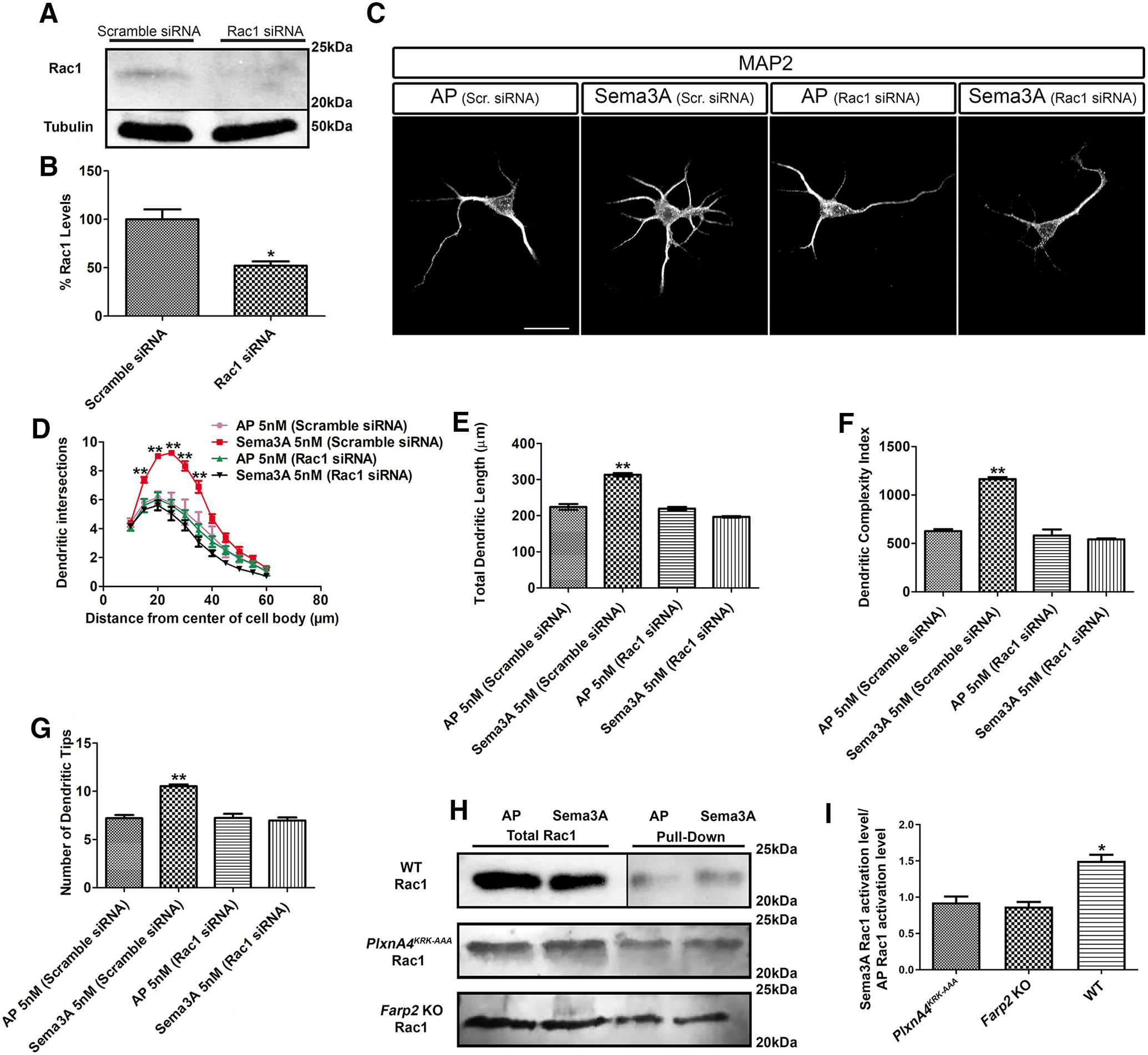
Sema3A increases Rac activation in cortical neurons and requires the presence of Rac1 for dendrite elaboration. ***A***, Rac1-specific siRNA decrease Rac1 total protein levels. **B**, Quantification of Rac1 levels from three independent experiments. *p < 0.05 (Student’s *t* test). ***C***, Representative confocal micrographs of dissociated primary cortical neurons obtained from WT E13.5 embryos transfected either with scramble siRNA or Rac1-specific siRNA. ***D***, Sholl analysis of dendritic intersections. ***E***, Total dendritic length. ***F***, DCI. ***G***, Number of dendritic tips for all treatment conditions described above. Data are mean ± SEM from three independent cultures. **p < 0.01 (two-way ANOVA with *post hoc* Tukey test). Scale bars: ***C***, 25 μm. ***H***, Pulldown assay showing the effect of 30 min stimulation of Sema3A on Rac1 activation in WT neurons, in *PlxnA4^KRK-AAA^*, and in *Farp2* KO neurons. ***I***, Quantification of Rac1-GTP fold change from three independent experiments. *p < 0.05 (one-way ANOVA).

## Discussion

How a finite number of molecular cues can sculpt the precise neuronal connections of complex circuits and induce diverse cellular responses is a fundamental question in neural development. Here, we show that Sema3A signals through the PlxnA4 receptor by selectively engaging the cytoplasmic KRK motif to specifically promote cortical neuron basal dendrite elaboration, but not CNS or PNS axon guidance. Although we cannot completely rule out the involvement of another coreceptor, which interacts with PlxnA4 in a KRK-dependent manner, currently there is no evidence supporting this notion. Our work supports the hypothesis that this KRK sequence is a signaling motif. First, FARP2 interacts with PlxnA4 in a KRK-dependent manner (Mlechkovich et al., 2014). Second, *PlxnA4^KRK-AAA^* and *Farp2* interact genetically *in vivo.* Additionally, this PlxnA4-FARP2 pathway requires the downstream target of FARP2, the small GTPase Rac1, to be active in cortical neurons but not in DRG sensory axons. Together, our results uncovered, for the first time, the selective Sema3A pathway through the KRK motif of the PlxnA4 receptor that activates FARP2-Rac1 signaling to control dendrite morphogenesis specifically in layer 5 cortical neurons (Fig. 10). In agreement with our work, a recent study suggested a functional divergence of the PlexinB motifs in the *Drosophila* olfactory circuit (Guajardo et al., 2019). Unlike our study, the downstream elements to these PlexinB motifs remain elusive.

**Figure 10.**
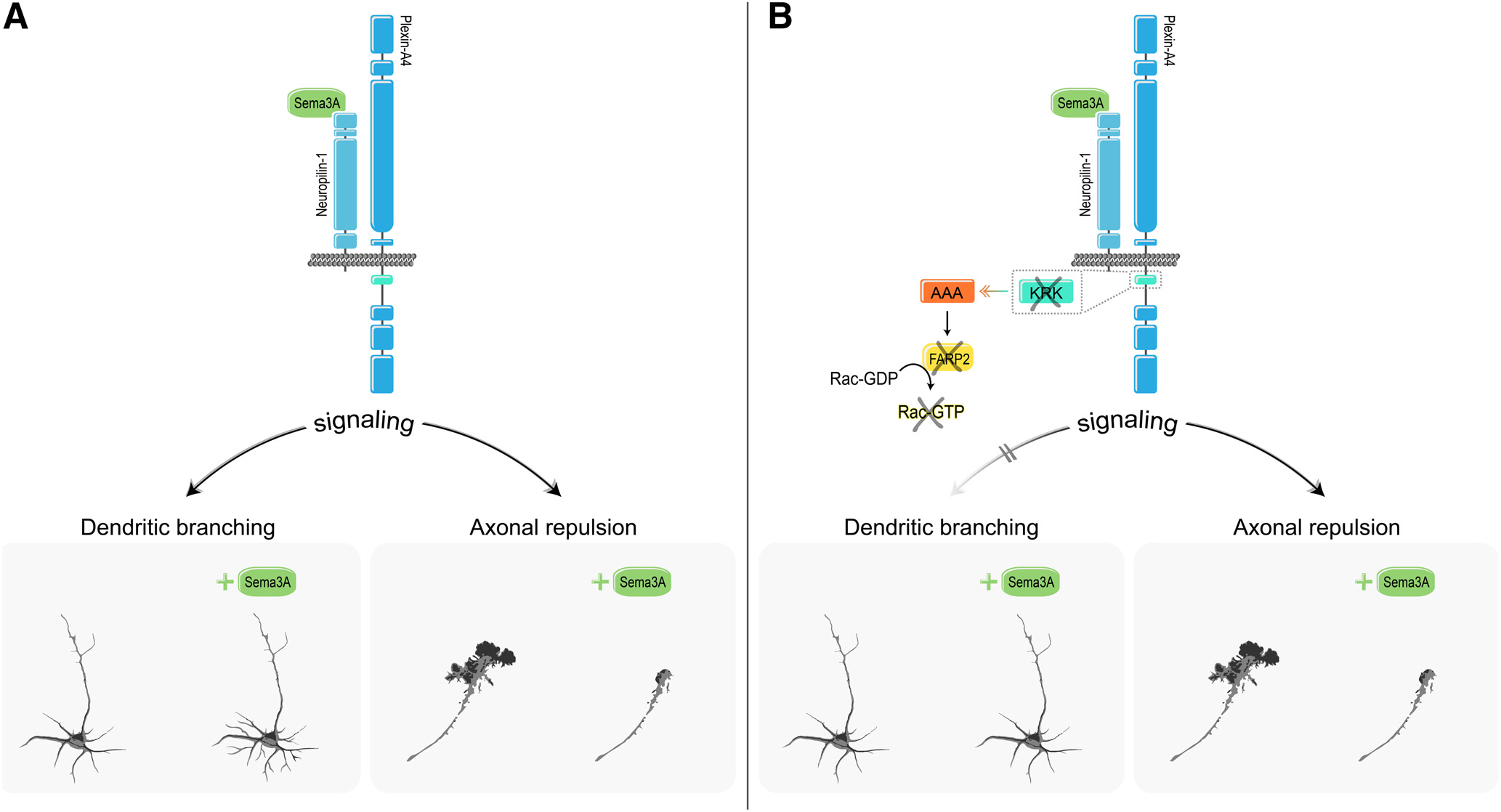
Differential requirement for the KRK motif and its downstream signaling effector FARP2 in Plexin-A4-mediated cellular responses. **A**, Sema3A-Nrp1/Plexin-A4 signaling promotes basal dendrite elaboration in deep-layer pyramidal cortical neurons on one hand and growth cone collapse and axonal repulsion on the other. **B**, Substitution of the KRK motif of Plexin-A4 to AAA, ablation of the Plexin-A4-binding effector, the Rac1 GEF FARP2, or inhibition of Rac1-specifically abrogate dendrite elaboration but not growth cone collapse and axonal repulsion.

### PlxnA4-FARP2-Rac1 pathway is required for dendrite morphogenesis but not for axon guidance

Our *in vitro* and *in vivo* analyses of the *PlxnA4^KRK-AAA^* mouse agree with our previous overexpression studies of PlxnA4 mutants in cultured neurons (Mlechkovich et al., 2014). While our previous study showed that Sema3A-mediated axon growth cone collapse is disrupted by *Farp2* siRNA knockdown in DRG neurons, our current results using the *Farp2* KO mouse contradict the *in vitro* knockdown findings. The basis for this difference may be due to off-target effects of the siRNA or compensatory mechanisms in the *Farp2* KO DRGs. Another study argued that Sema3A signaling mediates the dissociation of FARP2 from PlexinA1, resulting in the activation of Rac1 by FARP2 in axon repulsion assays of chick DRG neurons (Toyofuku et al., 2005). We have not found any evidence that this pathway is required for DRG axonal repulsion *in vitro* or for any repulsive guidance events of different neuronal types *in vivo*. Although there might be species differences between mice and chicken in Sema3A downstream signaling that may account for these discrepancies, our data support the model that this pathway is exclusively required for dendrite morphogenesis (Fig. 10).

While previous studies showed that Sema3A could increase Rac1 activity, they were mainly performed with either COS7 or HEK 293 cells (Turner et al., 2004; Toyofuku et al., 2005). We showed, for the first time, that Sema3A signaling in cortical neurons increases Rac1 activity. Another study using Rac1 siRNA in nonstimulated hippocampal neurons showed impairment in dendrite but not axon development (Gualdoni et al., 2007). In contrast, our data suggest that, in cortical neurons, Rac1 is needed for Sema3A-induced dendritic elaboration but not in the basal state. Furthermore, we corroborated our knockdown findings by inhibiting Rac1 activity with a pan-Rac inhibitor (EHT1864), which only perturbed Sema3A-mediated dendritic elaboration but not axonal growth cone collapse. While our results cannot rule out the contribution of other Rac isoforms (Rac2, Rac3, and RhoG), there are several reasons to support Rac1 as the favorite candidate. First, Rac2 expression has been shown to be restricted to hematopoietic cells (Roberts et al., 1999). Second, it has been proposed that Rac1 and Rac3 have opposing roles. While Rac1 is localized to the plasma membrane and cell protrusions, required for cell differentiation, neurite outgrowth, and involved in dendrites and spines development and maintenance (Luo, 2000; Gonzalez-Billault et al., 2012), Rac3 is localized in the perinuclear region, required for maintaining an undifferentiated state, and interferes with integrin-mediated cellmatrix adhesions (Corbetta et al., 2005; Hajdo-Milasinovic et al., 2007, 2009; Waters et al., 2008). The least studied is RhoG, which is expressed broadly (de Curtis, 2008) and previously shown to reduce dendrite complexity in hippocampal neurons (Franke et al., 2012; Schulz et al., 2016), rather than promoting dendritic arborization, as in our Rac1 results. Together, both *in vitro* and *in vivo* results support PlxnA4^KRK^-FARP2-Rac1 as the main pathway in Sema3A-mediated signaling in cortical neuron dendrite morphogenesis.

How is the specificity of the PlxnA4-FARP2-Rac1 pathway in cortical neuron dendritic morphogenesis achieved? One simple explanation is differential gene expression. However, all the components of this pathway are expressed both in cortical and DRG neurons during development (Mlechkovich et al., 2014). A recent single-cell analysis of DRG neurons at different developmental stages also detected the expression of FARP2, although at low levels, and expression of Rac1 (Sharma et al., 2020). Another possibility is that the KRK motif is inaccessible to FARP2 in DRG neurons. While beyond the scope of this study, this may be achieved by another FERM domain-containing protein, which is not required for signaling, or by differential lipid composition of the axons and dendrite membranes, as the KRK motif is located adjacent to the transmembrane domain.

### Sema3A signaling as a mechanism for promoting dendrite morphogenesis and remodeling in CNS neurons

Previous studies have identified the Src kinases (SFKs) as dowπ-stream effectors of Sema3A signaling in cortical basal dendrite elaboration (Sasaki et al., 2002; Morita et al., 2006). Recently, the protein tyrosine phosphatase *δ* (Ptp*δ*) was shown to interact with Nrp1 and to regulate the activity of SFKs in cortical neurons (Nakamura et al., 2017). However, these studies also demonstrated that the SFKs and Ptp*δ* control the growth cone collapse response mediated by Sema3A (Sasaki et al., 2002; Nakamura et al., 2017). Therefore, the SFKs do not confer functional specificity, but rather they may be considered as nonspecific signaling elements that are required for any type of Sema3A function.

While Sema3A can promote dendrite branching in dissociated hippocampal neurons (Fenstermaker et al., 2004), it also regulates the polarization of axon versus dendrite during hippocampal development (Shelly et al., 2011). Exogenous application of Sema3A suppressed undifferentiated neurites from differentiating into axons and promoted dendrite growth by elevating and reducing cGMP and cAMP levels, respectively. The ability of Sema3A to induce dendrite differentiation versus axon formation is recapitulated in *Xenopus* spinal neurons (Nishiyama et al., 2011). While the mechanism is unclear, Sema3A treatment of *Xenopus* spinal interneurons induced dendrite identity specification by simultaneous suppression of axon identity through regulation of cGMP signaling and voltage-sensitive Ca^2+^ channel Ca_V_2.3 expression. Activity-dependent Sema3A-induced dendrite branching is an interesting notion. Using an overexpression paradigm with tagged FARP1, Nrp1, and PlxnA1, another study showed the other member of the FERM domain containing RhoGEF FARP1 can associate with Nrp1/PlxnA1 complexes in HEK293 cells (Cheadle and Biederer, 2014). While Sema3A treatment of 21 DIV hippocampal neurons was unable to induce dendrite branching when FARP1 was knocked down in the presence of TTX, overexpression of PlxnA1 with TTX resulted in increased length of total dendritic branches, suggesting that the Semaphorin signaling pathway in hippocampal neurons is different from that in cortical neurons, which we show to be Sema3A-Nrp1/PlxnA4-FARP2-Rac1.

In conclusion, our study uncovers PlxnA4^KRK^/FARP2/Rac1 as a selective signaling pathway toward cortical pyramidal dendritic elaboration. Broadly, our results shed light on a longstanding question regarding the ability of multifunctional receptors to trigger diverse outputs, by demonstrating how distinct morphologic changes can be pinpointed to specific receptor signaling motifs and their downstream effectors.

